# Antimicrobial peptides signal need for sleep from peripheral wounds to the nervous system

**DOI:** 10.1101/2020.07.02.183806

**Authors:** Marina Sinner, Florentin Masurat, Jonathan Ewbank, Nathalie Pujol, Henrik Bringmann

## Abstract

Wounding triggers a protective innate immune response that includes the production of antimicrobial peptides and increased sleep. Little is known, however, about how peripheral wounds signal need for sleep to the nervous system. We found that during *C. elegans* larval molting, a tolloid/BMP-1-like protein promotes sleep through an epidermal innate immune pathway and the expression of more than a dozen antimicrobial peptide (AMP) genes. In the adult, epidermal injury activates innate immunity and turns up AMP production to trigger sleep. We show for one AMP, NLP-29, that it acts through the neuropeptide receptor NPR-12 in neurons that depolarize the sleep-active RIS neuron to induce sleep. Sleep in turn increases the chance of surviving injury. Thus, we found a novel mechanism by which peripheral wounds signal to the nervous system to increase protective sleep. Such a long-range somnogen signaling function of AMPs might also boost sleep in other animals including humans.

**Highlights:**

- Gain-of-function mutation in the tolloid/BMP-1-like NAS-38 protein increases sleep
- NAS-38 activates innate immunity pathways to ramp up STAT-dependent antimicrobial peptide (AMP) expression
- Wounding increases sleep through the innate immune response and AMPs
- Antimicrobial peptides are long-range somnogens that act through neuronal neuropeptide receptors to depolarize a sleep-active neuron
- Sleep increases the chance to survive injury

**Graphical Abstract:** 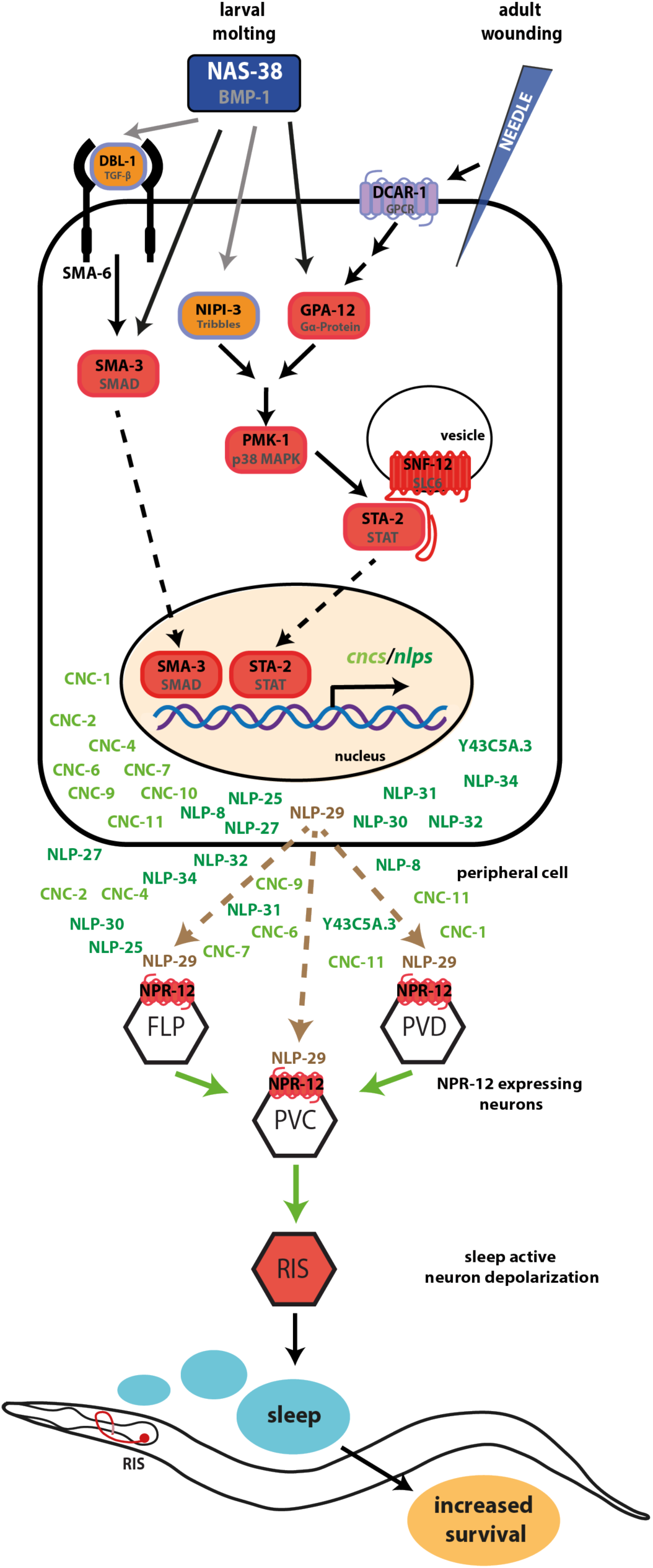

## Introduction

### Sleep supports the immune system

Following infection or wounding, the innate immune system is activated to isolate and destroy invading pathogens through the initiation of inflammatory processes. Part of the innate immune response in mammals is the so-called acute phase response, a systemic physiological state that creates a protective environment that includes fever and increased sleep. Specifically, non-REM sleep, which is believed to be restorative, is increased, whereas REM sleep, thought to be required for higher brain functions, is decreased [1, 2]. The amount of non-REM sleep correlates positively with the capacity of experimental animals to recover after infection [3]. More generally, proper sleep is needed to mount a normal immune response [4, 5]. Conversely, sleep loss is associated with impaired wound healing in animal models [6-8]. Among the effects of severe sleep deprivation are skin lesions and impaired adaptive as well as innate immunity, potentially leading to death by sepsis [9].

### Sleep is controlled by the immune system

As neural circuits control sleep, any impact of the immune system on sleep would be expected to be via the nervous system. From worms to humans, sleep is induced by sleep-active sleep-inducing neurons that express inhibitory neuropeptides and GABA. Physiological recordings have shown that these neurons depolarize specifically at the onset of sleep and optogenetic stimulation showed that they are sufficient to promote sleep. Ablation of these neurons impairs sleep, underscoring their importance in sleep induction [10-12]. Upstream circuits and molecular pathways control sleep-active neurons but little is known about the control of the activity of sleep-controlling circuits by the immune response. As part of the mammalian immune response, pro-inflammatory cytokines and prostaglandins appear to alter the excitability of neurons, but how altered excitability relates to the function of sleep circuits is unclear [1, 2, 4, 5, 13]. Sleep-controlling processes have mostly been studied in the brain. Peripheral tissues of the body, however, can also signal increased need for sleep. As examples, the circadian BMAL1 protein acts in muscle to control sleep amount, perhaps mediating the tiring effects of exercise [14], infection in peripheral tissues increases sleep to improve recuperation [5], and cellular stress can increase sleep through EGF signaling [15, 16]. However, little is known about the mechanisms by which peripheral tissues signal to the brain to control the activity of sleep-active neurons.

Antimicrobial peptides (AMPs) are also produced as a part of the innate immune response and can directly attack pathogens [17]. In *Drosophila*, an AMP called *nemuri* has been proposed to serve two functions. In peripheral tissues, its expression is increased upon infection and it may act directly in antimicrobial defense. In the brain, it is expressed and functions in neurons upstream of the dorsal fan shaped body, a region known to control sleep. Following sleep deprivation, *nemuri* expression is increased in the head and promotes sleep [18]. Thus, *nemuri* could be an AMP that has been co-opted to serve a somnogenic brain neurotransmitter function. In addition to sleep-controlling mechanisms in the brain, a long-range signaling mechanism must exist that signals from infected peripheral tissues to the brain. However, this mechanism remains unidentified and whether peripheral AMPs play a role in this process is unknown.

### *C. elegans* sleep

In *C. elegans*, sleep has been studied in the developing larva as well as in the adult [15, 19]. At the end of each larval cycle, *C. elegans* show sleep bouts during a developmental phase called lethargus, a period of time that coincides with synthesis of a new cuticle [20, 21]. Larval sleep bouts during lethargus are induced by the sleep-active neuron RIS, which is GABAergic and peptidergic and depolarizes during sleep bouts [22-24]. Optogenetic activation of the RIS neuron in a waking animal outside of lethargus leads to imminent sleep behavior. RIS is crucial for sleep, as worms with dysfunctional RIS have virtually no detectable sleep bouts. RIS inhibits arousal neurons and is itself inhibited by a strong, waking stimulus [22, 25]. Thus, RIS presents a *bona fide* sleep-active sleep-inducing neuron with strong similarities to mammalian sleep-active neurons.

### Innate immunity in *C. elegans*

*C. elegans* lacks adaptive immunity, but it possesses an innate immune system that responds to infection by a broad range of pathogens including viruses, bacteria, and fungi. Compared with vertebrates, innate immunity in the nematode is relatively simple as there are no specialized immune cells. This reduced complexity can be advantageous for dissecting its mechanisms [26, 27]. Besides pathogen infection, immune pathways in *C. elegans* can also be triggered by sterile injury of the epidermis [28-31]. Surveillance systems detect the presence of molecular patterns associated with physical damage, pathogens, and the cellular perturbation that they cause. Once such patterns are detected, intracellular signaling mechanisms are set off to induce the production of effectors that attack the pathogen or promote cell protection [32].

One of the key intracellular signaling pathways activated by infection in the worm consists of a series of kinase phosphorylation events that culminate in the phosphorylation and thus activation of a p38 MAP kinase [33]. p38 MAP kinase signaling plays a conserved role in promoting innate immune functions in both vertebrates as well as invertebrates [26, 34]. The p38 MAP kinase pathway can be triggered by several mechanisms, one of which involves activation of a damage-associated molecular pattern receptor (DAMP receptor) called DCAR-1 that then signals through a heteromeric G alpha protein called GPA-12 to activate a protein kinase C delta (TPA-1) [29]. An alternative activator during infection is the Tribbles homolog NIPI-3 [28]. PMK-1 triggers activation of a STAT transcription factor called STA-2 and AMP expression, a process that requires a SLC6 solute carrier protein called SNF-12 [28, 35, 36].

A second important pathway in *C. elegans* activated by pathogens is the TGF-β-SMAD pathway, which is activated by the extracellular ligand TGF-β/DBL-1, which is expressed on the outer surface of neurons and is released by proteolytic cleavage to act as a paracrine cue. It binds to the heteromeric TGF-β receptor formed by SMA-6 and DAF-4 on epidermal cells. Receptor activation during fungal infection then signals through the SMAD protein SMA-3 to boost AMP production in synergy with STA-2 [35, 37]. While to date in *C. elegans* TGF-β is only known to have a role in promoting innate immune responses, the role of this factor in other animals is more complex. In mammals, TGF-β is both a pro-inflammatory initiator of host responses that activates immune cells, but also is a potent suppressor of both innate and adaptive immunity [38].

Immunity pathways lead to the expression of protective proteins such as those regulating autophagy and the mitochondrial unfolded protein response as well as the production of molecules that directly attack intruding pathogens. These proteins include lysozymes that digest bacterial cell walls as well as AMPs that bind to and harm pathogens [27, 32, 34]. In *C. elegans*, infection and wounding cause the expression of many *cnc* and *nlp* genes through *sta-2*-dependent innate immunity acting cell autonomously in the epidermis [35]. These peptides are phylogenetically distinct from neuropeptides [39, 40], and all of these peptides are expressed in the epidermis, some in the intestine, and only a few in neurons [41]. Their overexpression confers resistance to pathogens [37, 39], and one of these genes was shown to directly attack pathogens [42]. Thus, this group of *nlp* and *cnc* genes is presumed to act as AMPs.

Here we aimed to find regulators of sleep in *C. elegans* through a genetic screen. We found a gain-of-function mutation in the tolloid/BMP-1-like *nas-38* gene, which caused substantially increased sleep during larval molting. NAS-38 acts through p38 MAP kinase and SMAD pathways to mobilize a STAT-transcription factor to produce a large battery of AMPs. In the adult, wounding activates this innate immune pathway and causes AMP expression to increase sleep. The AMPs in turn act on the nervous system to induce sleep through depolarization of the sleep-active RIS neuron. We show for one of the AMPs, NLP-29, that it acts on the nervous system via the neuropeptide receptor NPR-12, which is expressed in circuits that are upstream of RIS. Increased sleep promotes survival of injury demonstrating the biological significance of innate immunity-induced sleep. We thus delineate a novel cross-tissue signaling mechanism by which a peripheral tissue signals the need to sleep to the nervous system via AMPs.

## Results

### A genetic screen for increased larval sleep identifies a dominant mutation in the tolloid/BMP-1-like *nas-38* gene

In an attempt to identify novel genetic pathways controlling sleep in *C. elegans*, we conducted a large-scale reverse genetic screen for mutants that display increased larval sleep. We obtained a mutant strain for each gene for which a mutation was available at the Caenorhabditis Genetics Center. Out of about 4,500 mutant strains, 155 displayed increased behavioral quiescence (Supplementary Table 1). One strain, RB2467, clearly stood out among the high quiescence strains because it contained an unusually high fraction of immobile larvae that was much higher than in all other strains. Also, most quiescent worms did not show small occasional movements. This strain contained a deletion in the *nas-38* gene (*ok3407*), which was previously generated as part of a high-throughput allele project [43]. To measure how sleep behavior was altered by *nas-38(ok3407)*, after backcrossing this allele with wild type N2 worms, we quantified developmental time, lethargus length, and sleep behavior in mutants (Figure 1A-E). For this analysis, L1 larvae were cultured inside microfluidic chambers made from hydrogel and their development and behavior was imaged using DIC time-lapse microscopy [44]. Lethargus was identified as the continuous phase of pumping cessation prior to shedding of the cuticle and sleeping behavior was extracted by quantifying bouts of locomotion quiescence [21] (Figure 1A-B). *nas-38(ok3407)* L1 larvae developed with an almost normal speed until the onset of lethargus (time from hatching until the beginning of lethargus was 10.9 h on average in wild type and 12.1 h in *nas-38(ok3407)*, Figure 1C). The non-feeding phase, however, was specifically and dramatically extended more than twofold in *nas-38(ok3407)* (wild type and *nas-38(ok3407)* lethargus lengths were 1.7 and 3.6 h, respectively, Figure 1C-D. Also, the fraction of time spent in sleep was doubled (wild type was 23% and *nas-38(ok3407)* was 48%) and total sleep time during lethargus was about four-fold compared with the wild type (Figure 1E).

**Figure 1.**
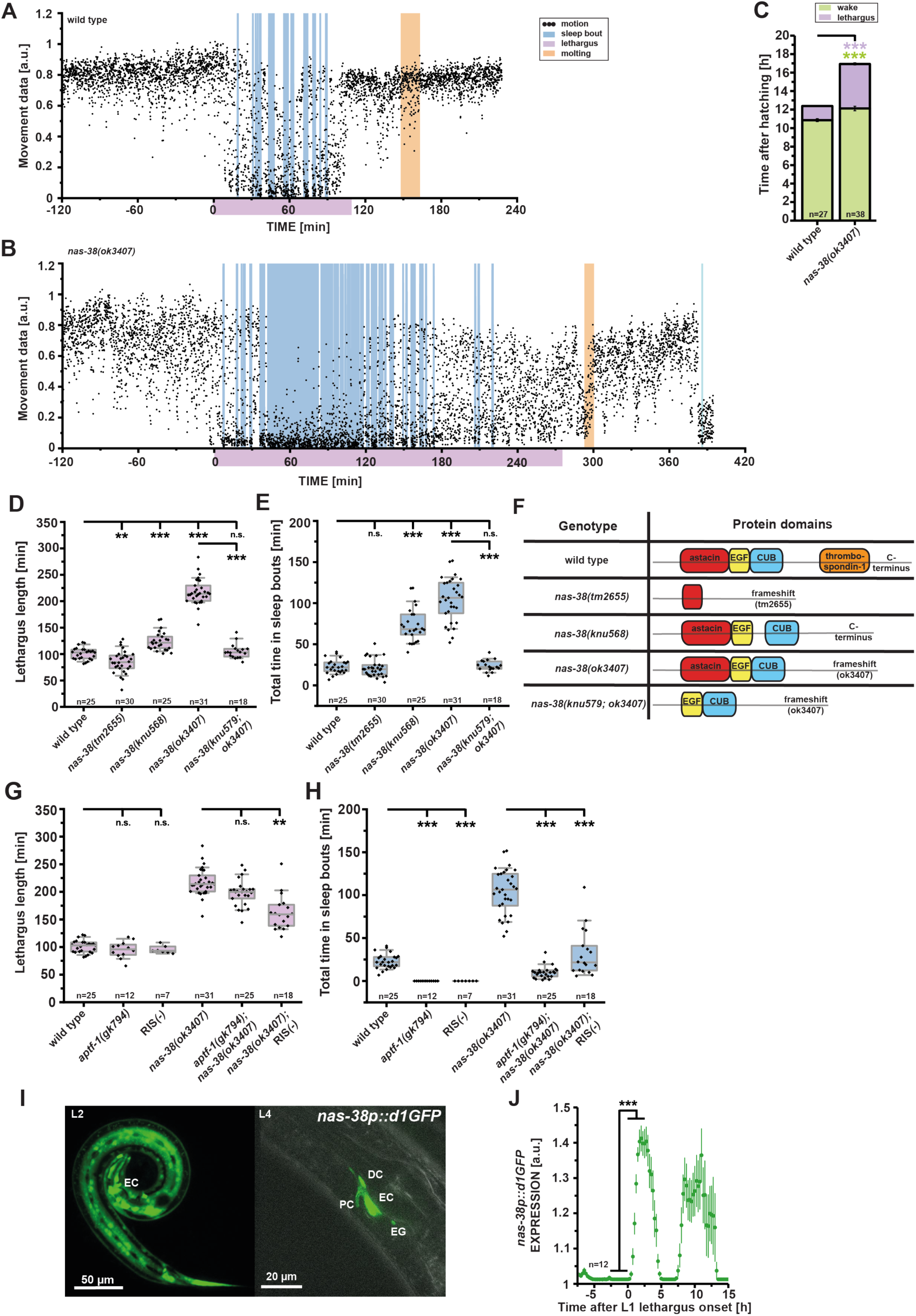
*nas-38* increases sleep during lethargus through the RIS neuron. (**A-B**) Image subtraction data of whole-body movement for one representative wild type and one *nas-38(ok3407)* worm before, during and after lethargus. In wild type animals, sleep bouts are only detectable during lethargus. (**C**) When measuring the total developmental time in L1, the lethargus length is specifically prolonged in the *nas-38(ok3407)* mutant. The time spent in the period before lethargus is increased by only 11.1 % on average but the mean lethargus is prolonged by 220.0 %, shown are mean values ± SEM, *** denotes p ≤ 0.001, two sample t-test. (**D**) Lethargus duration of different *nas-38* alleles. The loss-of-function (lf) allele *nas-38(tm2655)* shortens lethargus duration by 15.2 %. *nas-38(knu568)* increased only slightly by 23.4 %, *nas-38(ok3407)* prolongs lethargus by 115.7 %. ** denotes p ≤ 0.01, *** denotes p ≤ 0.001, two sample t-test. (**E**) Total time spent in sleep bouts in different *nas-38* alleles. *nas-38(knu568)* increased sleep by 219.0 %, *nas-38(ok3407)* increased sleep by 355.2 % compared to the wild type. This effect can be completely suppressed by a deletion of the astacin domain (77.3 % mean reduction of sleep compared to *nas-38(ok3407)*, no significant difference in *nas-38(knu579ok3407)* compared to wild types). *** denotes p ≤ 0.001, two-sample t-test. (**F**) Protein structures of different *nas-38* alleles that were used in this project. Frameshift mutation deletes the thrombospondin-1 domain and the C-terminus in *ok3407. nas-38(tm2655)* lacks most of the functional protein, including the astacin domain. *nas-38(knu568)* specifically lacks the thrombospondin-1 domain. *nas-38(knu579ok3407)* deletes the astacin domain in *ok3407*. (**G**) RIS(-) decreased lethargus duration of *nas-38(ok3407)* by 25.3 %. Data are compared against the same wild type and *nas-38(ok3407)* data as in (**E**). ** denotes p ≤ 0.01, two-sample t-test. (**H**) In wild types and in *nas-38(ok3407)* mutants, loss of RIS reduced sleep time during lethargus. *aptf-1(gk794)* and RIS(-) suppressed sleep. In *nas-38(ok3407), aptf-1(gk794)* reduced the sleep amount by 90.3 % and RIS(-) by 69.1 %. The same wild type and *nas-38(ok3407)* sleep data as in (**D**) are plotted in this panel. *** denotes p ≤ 0.001, two-sample t-test. (**I**) Left panel: L2 larva expressing *nas-38::GFP* during lethargus inside a microchamber. GFP is mainly expressed in the epidermis except the seam cells. EC indicates the excretory cell. Right panel: immobilized L4 larva during lethargus. Shown is the merged image of the brightfield and GFP channel. Pore cell (PC), duct cell (DC) and the excretory gland cells (EG). (**J**) *nas-38* expression oscillates with lethargus. During lethargus it increases by 20.0 %, ***denotes p ≤ 0.001, paired Wilcoxon rank test, and drops afterwards. Boxplots show individual data points, the box represents the 25 % – 75 % range, the whiskers cover the 10 % - 90 % range, the thin gray line is the median. The same wild type and *nas-38(ok3407)* data as in figures 4A-D are shown in every panel.

To test whether the sleep phenotype was recessive or dominant, we generated and tested worms that were heterozygous for *nas-38(ok3407)*. The heterozygous mutation of *nas-38(ok3407)* showed a modest increase in lethargus length and total sleep time that was much lower compared with the homozygous mutant, suggesting that *nas-38(ok3407)* is semi-dominant (Figure S1A-B, lethargus duration was 28.1 % longer in comparison to wild type, but 40.6 % shorter compared to the homozygous *ok3407* allele, sleep time in heterozygous *nas-38(ok3407)* was 69.9 % less compared to the homozygous *ok3407* allele). *nas-38(ok3407)* causes a deletion of the C-terminus region in the protein and a frameshift that leads to an artificial C-terminus. To test whether the C-terminal deletion of *nas-38* or the artificial C-terminus caused the dominant sleep phenotype, we rebuilt a clean deletion of the C-terminus by introducing a stop codon at the same position where *nas-38(ok3407)* is truncated using CRISPR/Cas9 [45] (Figure S1C). This clean truncation, *nas-38(syb293)*, completely recapitulated the sleep phenotype of *nas-38(ok3407*), confirming that the phenotype is indeed caused by removal of the C-terminal part of *nas-38* that includes a thrombospondin (TSP1) domain (Figure S1A-B) (334.2 % increased sleep compared to wild type, 116.0 % increased lethargus length).

To assess the loss-of-function phenotype of *nas-38*, we analyzed sleep in a deletion mutant, *nas-38(tm2655)* that removes the majority of the gene product and likely presents a molecular null. *nas-38(tm2655)* showed a modest reduction of lethargus time and sleep behavior (Figure 1D-E, 15.2 % lethargus decrease, n.s. sleep difference). Together these results indicate that *nas-38(ok3407)* and *nas-38(syb293)* are strong gain-of-function alleles that dramatically increase lethargus length and sleeping behavior. Thus, *nas-38* promotes lethargus duration and sleeping behavior.

NAS-38 belongs to the family of astacin zinc metalloproteases that are found in all species investigated ranging from bacteria to humans. Astacin metalloproteases play diverse roles in extracellular matrix processing as well as in extracellular signaling pathways. The vertebrate bone morphogenic protein 1 (BMP-1) acts as a procollagen C-protease to induce bone formation and also plays a role in activating TGF-β signaling [46]. A BMP-1/Tolloid homolog called BMP-1 exists in *C. elegans.* BMP-1 belongs to a group of 40 paralogous genes called the Nematode Astacins (NAS). BMP-1 consists of a signal peptide that is followed by the Astacin protease domain and several CUB and EGF repeats. The structure of NAS-38 is similar to BMP-1 but has a reduced number of CUB and EGF domains and has an additional TSP1 domain (Figure 1F) [46].

To identify structures critical for NAS-38 function, we used CRISPR/Cas9-edited *nas-38* alleles. Because the *nas-38(ok3407)* deletion includes the TSP1 domain, we tested whether it is the absence of this domain that caused the gain of function phenotype. We generated a specific in-frame deletion of the TSP1 domain, *nas-38(knu568)*. While lethargus duration was increased only slightly, sleep was strongly increased in *nas-38(knu568)*, showing that increased sleep is caused mostly by TSP1 absence, suggesting that the TSP1 domain normally represses NAS-38 activity (58% higher fraction of time spent in sleep, 219.0 % more total sleep, 23.4 % longer lethargus compared to wild type, Figure 1D-E). To test whether *nas-38*(*ok3407)* acts through its astacin domain, we engineered an in-frame deletion of this domain in the gain of function allele (*nas-38(ok3407knu579)*). Astacin domain deletion completely suppressed the increased sleep and lethargus length phenotype of *nas-38(ok3407)*, suggesting that protease activity is essential for NAS-38 function (sleep amount and lethargus length were not statistically significantly different to wild type worms, but compared to *nas-38(ok3407)*, sleep was shortened by 77.3 % and lethargus was reduced by 51.6 %, Figure 1D-E). Together, these results suggest an inhibitory function for the TSP1 domain in NAS-38 and indicate that the astacin protease domain is required for NAS-38 functions. The lethargus and sleep phenotypes can be influenced to various extents and are caused by an absence of the complete C-terminus or only the TSP1 domain, respectively.

### NAS-38 is expressed in peripheral tissues and increases sleep through the RIS neuron

Sleep bouts during lethargus are induced by the sleep-active RIS neuron [22, 25]. To test whether sleep bouts in *nas-38(ok3407)* depend on RIS, we used a genetic inactivation of RIS (RIS(-), caused by expression of *egl-1*, (Wu et al., 2018)) as well as a molecular null mutation in *aptf-1*, which almost completely lacks sleep bouts. APTF-1 is a transcription factor required for the expression of sleep-inducing neuropeptides in RIS [22, 25]. We measured the effects of RIS manipulation on lethargus length and sleep amount in *nas-38(ok3407)*. In the *nas-38(ok3407)* mutant background, RIS impairment reduced the lethargus length by 25.3 % (Figure 1G). *aptf-1(gk794)* reduced the total sleep time in *nas-38(ok3407)* mutants by 90.3 %, and RIS(-) by 69.1 %, (Figure 1H, the effects of RIS(-) are variable due to silencing of the transgene, which might explain the weaker effect in RIS(-)). These results indicate that *nas-38(ok3407)* increases sleep mostly by acting through RIS.

We generated a transcriptional (promoter) fusion with a destabilized fluorescent reporter (d1GFP) [47] and followed *nas-38* expression across larval development and the sleep-wake cycle using fluorescence microscopy time-lapse imaging of worms cultured inside microfluidic devices. *nas-38* was expressed in the epidermis but also in the excretory canal cell, duct cell, pore cell and excretory gland cell but not in the RIS neuron (Figure 1I). Expression was first detectable in the embryo. During L1 lethargus, expression strongly increased and decreased again after the end of lethargus during the L2 stage and rose again during L2 lethargus (Figure 1J). Thus, consistent with high-throughput expression data, *nas-38* is expressed in epithelial cells in an oscillating expression pattern with an increase during lethargus [48, 49].

### *nas-38* promotes expression of immune response genes

To find out how *nas-38* controls lethargus length and sleeping behavior as well as to identify the downstream effectors through which NAS-38 signals, we analyzed the transcriptomes of *nas-38* mutants during lethargus. We manually isolated wild type, *nas-38(tm2655)* loss-of-function, and *nas-38(ok3407)* gain-of-function worms during L4 lethargus based on the absence of pharyngeal pumping. The worms were then subjected to RNAseq and differentially expressed genes were extracted. The transcriptome of *nas-38(tm2655)* showed only mild changes compared with the wild type. By contrast, the transcriptome of *nas-38(ok3407)* was massively changed with hundreds of genes changing their expression compared with the wild type (Figure 2A). The strength of the gene expression changes in the mutants thus matched the behavioral phenotypes for the two *nas-38* mutations, with modest alterations seen in the loss-of-function mutation and massive changes observed in the gain-of-function mutation.

**Figure 2.**
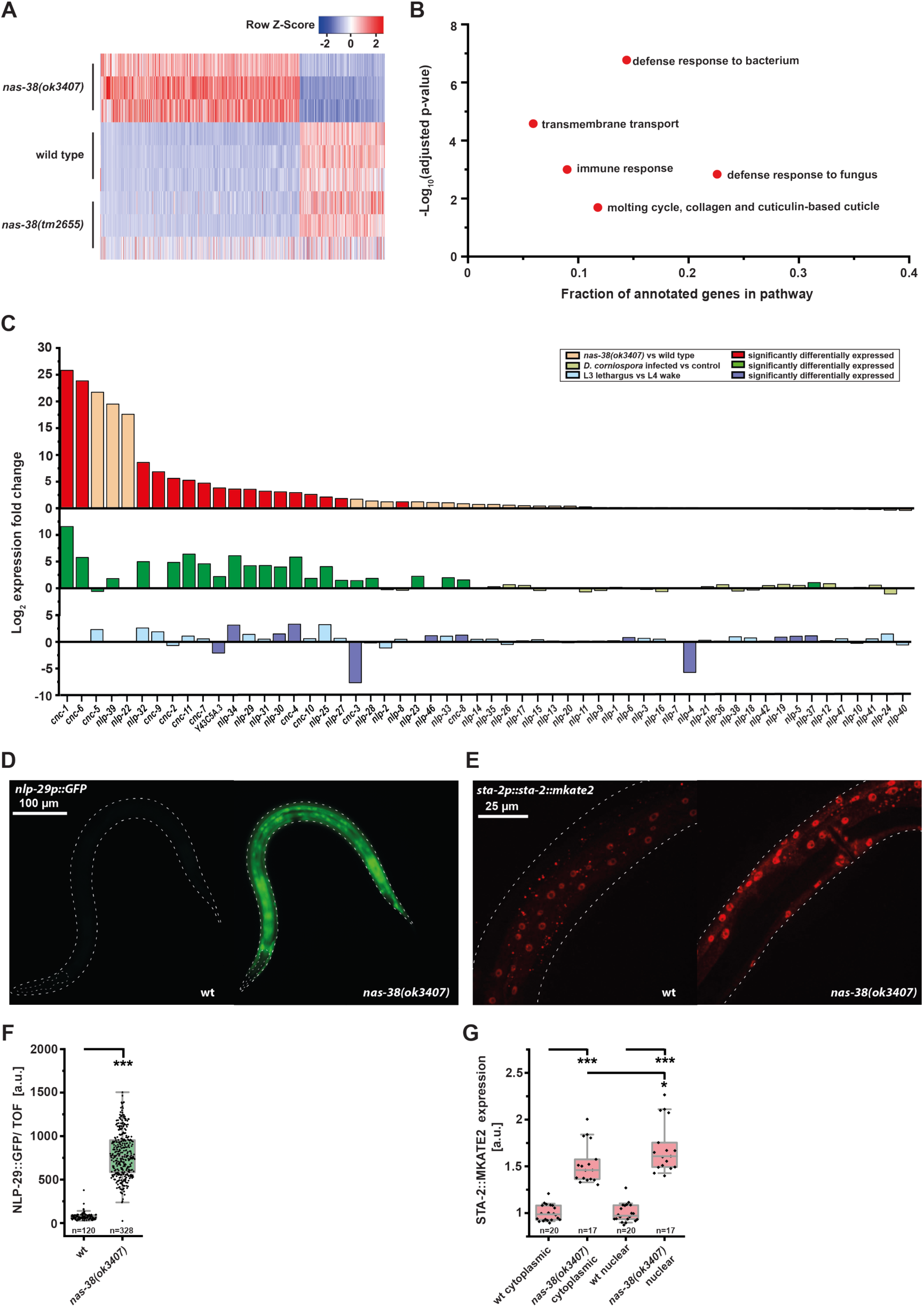
NAS-38 induces innate immune response gene expression. (**A**) Heatmap of the statistically significant transcriptional changes in *nas-38(ok3407)* and *nas-38(tm2655)* compared to wild type. (**B**) Gene set enrichment analysis of differentially expressed genes reveals similarities with immunity and defense responses as well as molting. (**C**) Comparison of differential expression of *nlps* and *cncs* in *nas-38(ok3407)*, infected worms and wild type larvae during lethargus. Differentially expressed genes of the infection and lethargus transcriptomes were compared with the *nas-38(ok3407)* transcriptome by negative binomial testing. Light shading indicates a non-significant up or down regulation, dark shading indicates a significant expression fold change. *nas-38gf*-induced *nlp* and *cnc* upregulation is most similar to infected worms. (**D**) Expression of *nlp-29p::GFP* is strongly increased in *nas-38(ok3407)* L4 larvae. Expression in the wild type is visible only under conditions that caused saturation of the signal in *nas-38(ok3407)*. (**E**) A STA-2::mKate2 fusion protein expressed under the endogenous *sta-2* promoter revealed an increase in expression as well as nuclear localization of STA-2. (**F**) Fluorescence quantification reveals an upregulation of *nlp-29p::GFP* by 966.6 % in a mixed population in the *nas-38(ok3407)* mutant. (**G**) Cytoplasmic and nuclear STA-2::mKate2 quantification. Data were normalized to wild type levels. In *nas-38(ok3407)*, cytoplasmic STA-2 increased 1.5 fold and nuclear STA-2 increased 1.7 fold. * denotes p ≤ 0.05, two-sample t-test. The boxplots show individual data points, the box represents the 25 % – 75 % range, the whiskers cover the 10 % - 90 % range, the thin gray line is the median.

To identify key biological processes that are affected by *nas-38(ok3407)*, we performed tissue and gene enrichment analyses. Tissue enrichment analysis showed that the differentially expressed genes in *nas-38(ok3407)* were most strongly associated with secretion, which is consistent with the expression pattern of *nas-38* in excretory tissues (Figure S2). Gene enrichment analysis showed that the genes that were upregulated by *nas-38(ok3407)* were most strongly associated with formation of the cuticle and with innate immunity responses (Figure 2B). The innate immune response involves the production of AMPs such as NLPs (neuropeptide like proteins) and CNCs (caenacins) [28, 40]. AMPs of the *nlp* and *cnc* gene families changed their expression up to several hundred fold (Figure 2C).

Epidermal AMPs are synthesized during infection but also after sterile wounding and in specific cuticular collagen mutants [28, 50]. Epidermal AMPs are also expressed specifically during molting [49, 51-53]. Hypothetically, *nas-38(ok3407)* could thus either act by increasing molting-related AMP gene expression or it could trigger expression of AMPs required for immune response. We therefore tested whether the set of *nlp* and *cnc* genes that were overexpressed in *nas-38(ok3407)* were more similar to those AMPs that are induced by the immune response or to those that are rhythmically expressed during lethargus. The significantly upregulated hits of the *nas-38(ok3407)* transcriptome corresponded to 15 out of 17 cases with significantly upregulated AMPs after infection with *Drechmeria coniospora* and only in 3 out of 17 cases with the transcriptome data obtained from worms in lethargus (Figure 2C). This suggested that *nas-38(ok3407)* generates an AMP response that is mostly similar to the innate immune response.

To validate the transcriptome changes in *nas-38(ok3407)*, we looked at reporter gene expression of NLP-29. NLP-29 has been shown to be representative for the AMP peptide cluster and the promoter fusion transgene recapitulates the increased expression in the epidermis after infection and wounding [28, 42]. Whereas expression during lethargus from the *nlp-29* promoter was barely detectable in wild type animals, this promotor expressed strongly in the epidermis in *nas-38(ok3407)* (Figure 2D).

Expression of *sta-2* increased 4.3-fold in *nas-38(ok3407)* in the transcriptome. Upon activation, STA-2 is presumed to translocate into the nucleus [35]. We thus tested whether *nas-38(ok3407)* affects not only the expression level of STA-2 but also impacts its translocation into the nucleus. We followed the localization of a translational reporter fusion of STA-2 in epidermal cells during lethargus and quantified the fraction of STA-2 that was localized to the nucleus. The translational reporter for STA-2 was enriched in the nuclei of epidermal cells but was also found in the cytoplasm. The reporter was indeed expressed at 1.5-fold higher levels in the cytoplasm in the *nas-38(ok3407)* background. In addition, *nas-38(ok3407)* led to an increase of the nuclear fraction of STA-2 (Figure 2E-F). In summary, *nas-38(ok3407)* led to massive transcriptional alterations, activation of the innate immune response, and increased expression of AMPs that are expressed in peripheral tissues.

### NAS-38 promotes sleep through innate immune response genes

*nas-38(ok3407)* induces a massive immune-like peripheral transcriptional response during lethargus. We thus hypothesized that innate immune genes are responsible for the increased lethargus length and induce sleep behavior. To identify downstream effectors through which *nas-38* acts, we performed epistasis experiments with genes that are known to be involved in the epidermal immune response.

We focused on two conserved pathways, the first acting through a p38 MAP kinase and the second acting through DBL-1/TGF-β-SMAD signaling (Figure 3A). We crossed mutants of key genes of these pathways into our gain-of-function mutants *nas-38(knu568)* and *nas-38(ok3407)*, and quantified quiescence behavior. Our earlier analysis indicated that *nas-38(ok3407)* presents a more severe gain-of-function allele with the strongest increase in lethargus length, whereas *nas-38(knu568)* presents a more moderate gain-of-function allele in which sleep amount is specifically increased. Mutation of the most upstream components of the p38 MAP kinase pathway *dcar-1* (damage receptor) and *nipi-3* (tribbles) did not suppress increased sleep behavior of *nas-38(ok3407)* worms (Figure S3A and B). Mutations of *gpa-12* (G alpha protein), *pmk-1* (p38 map kinase), *snf-12* (solute carrier), and *sta-2* (STAT) that were further downstream in the pathway could, however, suppress increased sleep (lethargus length in *nas-38(ok3407)* was suppressed by 14.9 % in *gpa-12* mutants, by 17.5 % in *pmk-1* mutants, by 29.6 % in *sta-2* mutants and by 49.8 % in *snf-12* mutants, Figure 3B-C, S3A-C).

**Figure 3.**
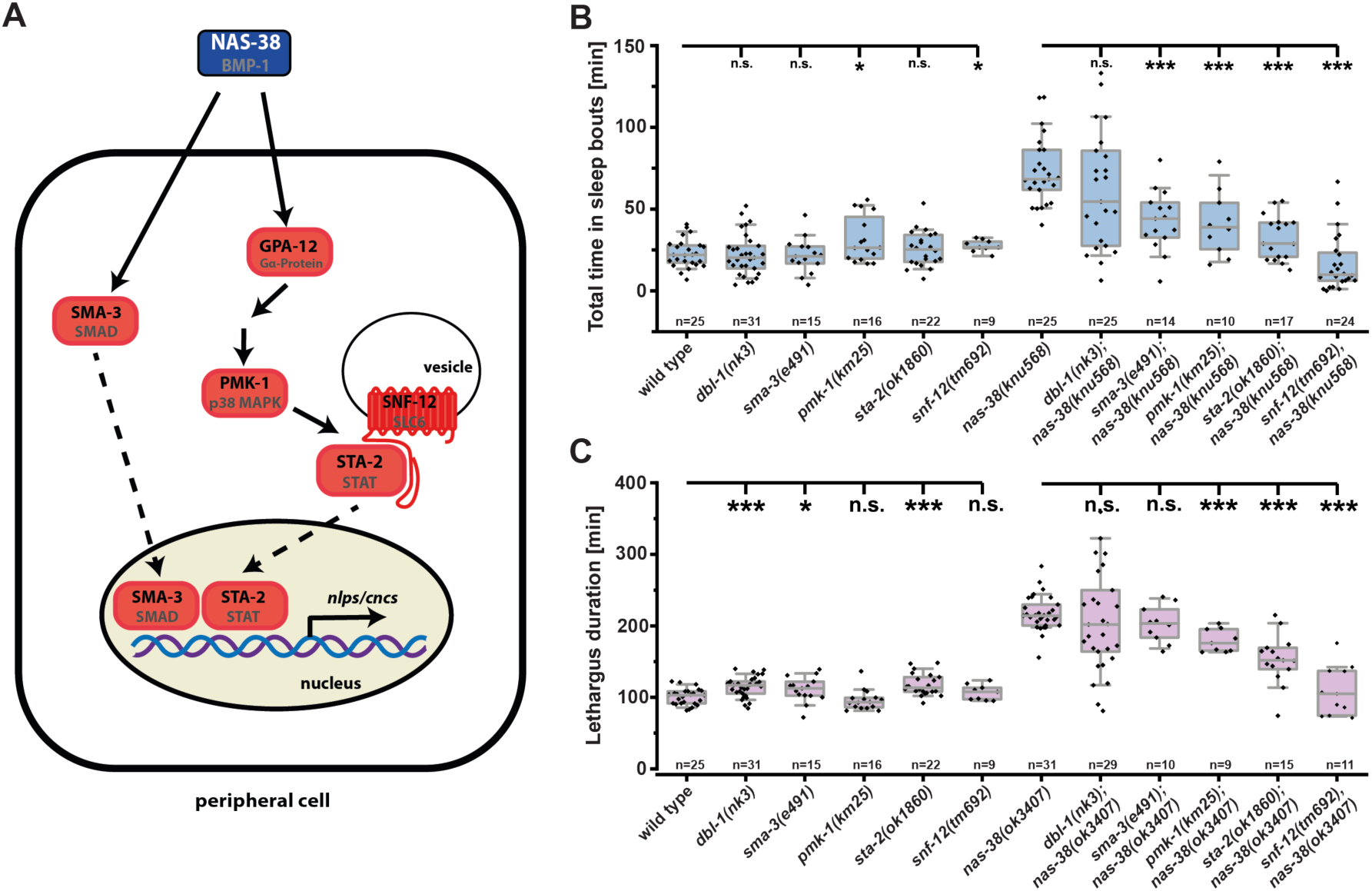
NAS-38 increases sleep and lethargus length through TGFβ/SMAD and p38 MAP-Kinase innate immune response pathways. (**A**) Cartoon summarizing the pathways involved in mediating increased sleep caused by *nas-38*. Proteins corresponding to mutants that suppressed the sleep phenotype of *nas-38* gain-of-function mutants are marked in red. (**B**) *sma-3(e491)* reduced sleep in *nas-38(knu568)* by 41.2 %, *pmk-1(km25)* by 43.7%, *sta-2(ok1860)* by 55.5 % and *snf-12(tm692)* by 77.2 %, * denotes p ≤ 0.05, *** denotes p ≤ 0.001, two sample t-test. (**C**) *pmk-1(km25)* decreased lethargus length in *nas-38(ok3407)* by 17.5 %, *sta-2(ok1860)* by 29.6 %, and *snf-12(tm692)* by 49.8 %. * denotes p ≤ 0.05, *** denotes p ≤ 0.001, two sample t-test. The boxplots show individual data points, the box represents the 25 % – 75 % range, the whiskers cover the 10 % - 90 % range, the thin gray line is the median.

For the TGF-β-SMAD signaling pathway, mutations of *dbl-1* (TGF-β ligand) and *sma-3* (SMAD) could not suppress increased quiescence in *nas-38(ok3407)* worms. However, a mutation in *sma-3* suppressed *nas-38(knu568)*-increased sleep by 41.2 %. Suppression of increased sleep was more clearly visible for all tested pathway mutants in *nas-38(knu568)*, perhaps because the sleep phenotypes is stronger and more specific and thus easier to suppress in this mutant (Figures 3B and S3C). The increased lethargus length in *nas-38(ok3407)* mutants was suppressed by the same genes that suppressed increased sleep in *nas-38(knu568)*, but lethargus length suppression was stronger in the *nas-38(ok3407)* allele that had the highest increase in lethargus length (Figure 1D). Thus, to increase sleep behavior, *nas-38* gain-of-function mutations act through at least two pathways, the highly conserved wounding and infection-responsive p38 MAP kinase and possibly also through a SMAD signaling pathway, both of which are known to activate SLC6/SNF-12 and STA-2 in the epidermis [35].

### NAS-38 increases epidermal antimicrobial peptide expression through innate immunity genes

Many *cnc* and *nlp* genes were upregulated during lethargus in *nas-38(ok3407)* (*cnc-1, cnc-2, cnc-4, cnc-6, cnc-7, cnc-9, cnc-10, cnc-11, nlp-8, nlp-25, nlp-27, nlp-29, nlp-30, nlp-31, nlp-32, nlp-34, Y43C5A.3*). Most of these genes are known to be regulated by SNF-12, STA-2 and/or SMA-3 in response to wounding and infection and are presumed to act as AMPs [28, 35, 37]. We hence wanted to test the hypothesis that *nas-38* controls AMP expression through the immune response pathway. To study the expression of the AMPs, we looked at a transgenic reporter line that expresses GFP under the control of the *nlp-29* promoter [28]. We cultured L1 larvae in microfluidic devices, took time-lapse fluorescence movies, and quantified *nlp-29* expression. *nlp-29* expression oscillated with the developmental cycle and increased strongly during lethargus, consistent with high-throughput transcriptome data [49, 51] (Figure 4A). We next looked at the effects of *nas-38(ok3407), nas-38(knu568)* and *nas-38(tm2655)* on *nlp-29* expression. In *nas-38(ok3407)*, expression of *nlp-29* was strongly increased, in *nas-38(knu568)* expression was moderately increased and in *nas-38(tm2655)* the expression was decreased by about half (Figure 4A). Thus, the level of *nlp-29* expression correlated with the amount of sleep. We next assayed the suppression of overexpression of *nlp-29* in *nas-38(ok3407)* by components of the innate immune response pathway that we had tested for quiescence suppression before. The expression of *nlp-29* was suppressed by genes of the p38 MAP kinase pathway and, to a lesser degree, by genes of the DBL-1/TGF-β-SMAD signaling pathway (*gpa-12(pk322)*: 71.3 %, *sta-2(ok1860)*: 97.5 %, *snf-12(tm692)*: 81.0 % (Figure 4B-E); *nipi-3(fr4):* 51.8%, and *pmk-1(km25):* 54.2 %, *dbl-1(nk3)*: 28.5 %, *sma-3(e491)*: 33.4 % (Figure S4A-G)). Thus, *nas-38* acts through innate immune response genes to increase *nlp-29* expression in the epidermis. Overall, high expression of *nlp-29* in mutants correlated with increased sleep behavior, and suppression of *nlp-29* expression was correlated with suppression of increased sleep behavior (Figure 4F).

**Figure 4.**
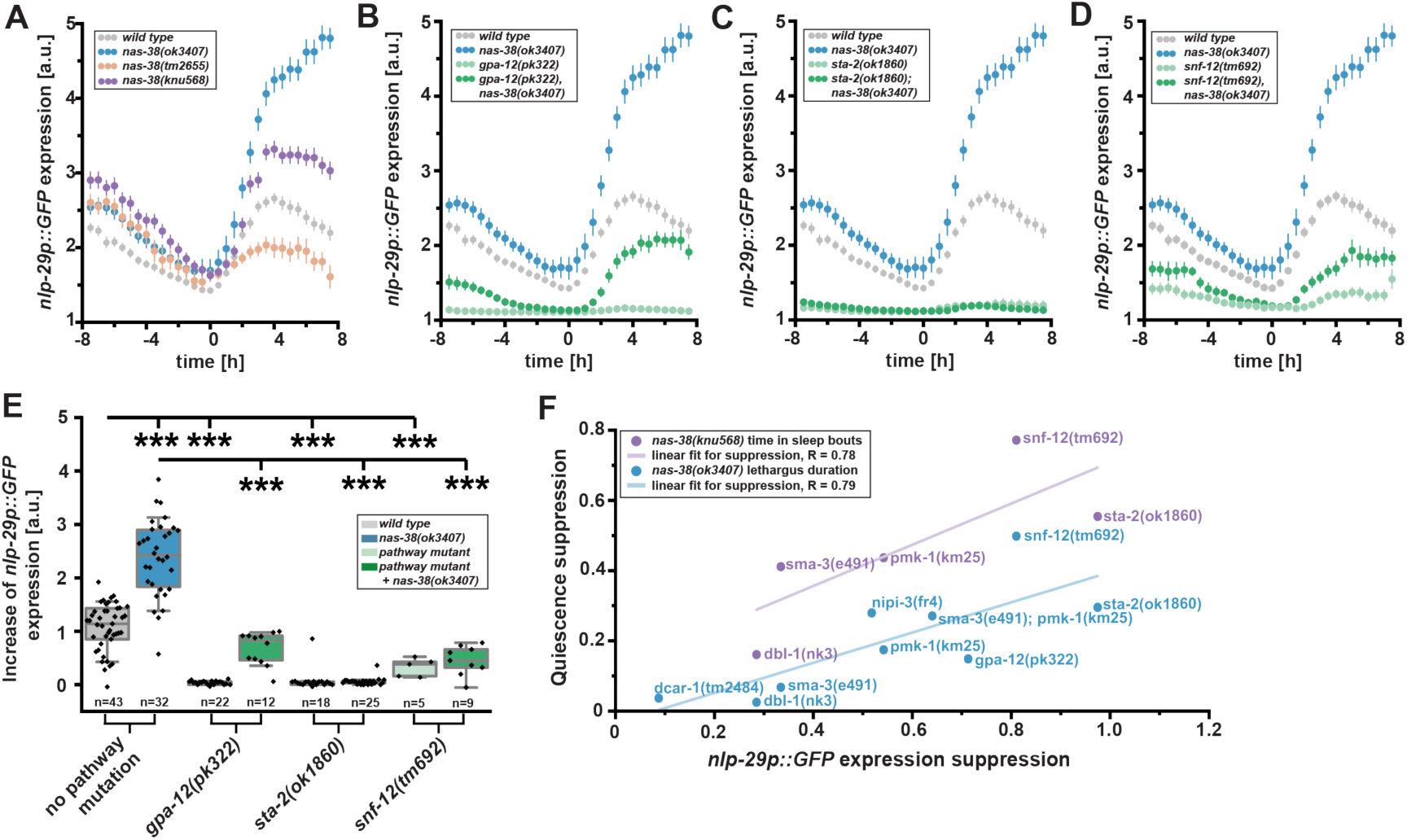
NAS-38 controls the expression of NLP-29 through innate immunity pathways. (**A**) – (**D**) *nlp-29p::GFP* intensity levels 8 h before and after lethargus onset. Wild type and *nas-38(ok3407)* data are repeatedly shown in every panel. (**A**) *nas-38(ok3407)* increased *nlp-29p::GFP* expression by 120.5 %, *nas-38(knu568)* increased *nlp-29p::GFP* expression by 45.1 % and the *nas-38(tm2655)* allele decreased *nlp-29p::GFP* expression by 69.5 % 4 h after lethargus onset. (**B**) – (**D**) Suppression of *nas-38(ok3407)*-induced *nlp-29p::GFP* expression in mutants of innate immune pathways. Wild type expression levels are shown in grey, expression levels in single mutants in light green and dark green in double (or triple) mutants with *nas-38(ok3407). gpa-12(pk322), sta-2(ok1860)* and *snf-12(tm692)* suppressed *nlp-29p::GFP* expression in *nas-38(ok3407)*. Shown are mean values ± SEM. (**E**) *nlp-29p::GFP* expression over the first 4 h after lethargus onset. *gpa-12(pk322)* decreased *nas-38(ok3407)*-induced *nlp-29p::GFP* expression by 71.3 %, *sta-2(ok1860)* by 97.5 % and *snf-12(tm692)* by 81.0 % ** denotes p ≤ 0.01, *** denotes p ≤ 0.001, two sample t-test. The boxplots show individual data points, the box represents the 25 % – 75 % range, the whiskers cover the 10 % - 90 % range, the thin gray line is the median. (**F**) Suppression of *nlp-29p::GFP* expression correlates with suppression of lethargus increase and sleep increase in *nas-38* gain-of-function alleles. The more a mutation suppressed the quiescence phenotype, the stronger it also suppressed *nlp-29p::GFP* expression.

### *nas-38* acts through AMPs to increase sleep

As sleep correlated with AMP expression, we hypothesized that *nas-38(ok3407)* increases sleep through AMPs. We crossed knockout alleles of an individual AMP, *nlp-29* and another neuropeptide-like gene that was overexpressed in *nas-38(ok3407), nlp-8*, into the *nas-38(ok3407)* mutant and analyzed lethargus duration and sleeping behavior using time lapse microscopy. Neither the deletion of *nlp-29* nor *nlp-8* could suppress increased sleep or lethargus duration in *nas-38(ok3407)* (Figure S5A and S5B). As the members of the *nlp* and *cnc* gene families are structurally highly similar [39, 40] they may act redundantly. To test this hypothesis, we created a strain using CRISPR/Cas9 that deleted 16 out of 17 *nlp* and *cnc* genes that were upregulated in the *nas-38(ok3407)* transcriptome. As four of the significantly upregulated *cnc* genes and five of the *nlp* genes are organized in gene clusters, we decided to delete those two genomic regions completely. This resulted in the knockout of a total of 19 *nlp* and *cnc* genes (the significantly upregulated *cnc-1, cnc-2, cnc-4, cnc-6, cnc-7, cnc-10, cnc-11, nlp-8, nlp-25, nlp-27, nlp-29, nlp-30, nlp-31, nlp-32, nlp-34* and *Y43C5A.3* as well as the additional cassette members *cnc-3, cnc-5* and *nlp-28*, which also were upregulated in the *nas-38(ok3407)* transcriptome, yet did not reach statistical significance).

The multi *nlp* and *cnc* deletion strain did not have sleep or lethargus defects on its own. However, *nlp* and *cnc* deletion suppressed increased sleep of *nas-38(ok3407)* by 27.6 % as well as lethargus duration by 17.0 % (Figures 5A-D). The suppression of increased sleep was especially prominent in the first part of the lethargus phase (Figure 5D). We also knocked down all the significantly upregulated *nlp* and *cnc* genes simultaneously by RNAi. For this experiment, we generated a feeding RNAi construct that combined the coding sequences of all *nlp* and *cnc* genes that were significantly upregulated in *nas-38(ok3407)*. The multi *nlp* and *cnc* RNAi knockdown significantly suppressed sleep behavior in *nas-38(ok3407)* mutants by 15.3 % and lethargus duration by 6.4 % (Figure S5C and S5D). Thus, *nlp* and *cnc* genes increase sleep and lethargus duration of *nas-38(ok3407)*. The partial *nas-38(ok3407)* suppression phenotype suggests that additional genes are involved in sleep induction. Additional potential AMP genes exist in *C. elegans* [39] that are up regulated by both *nas-38(ok3407)* mutant and epidermal infection, and thus perhaps even more AMPs might be involved in immunity-triggered sleep induction.

**Figure 5.**
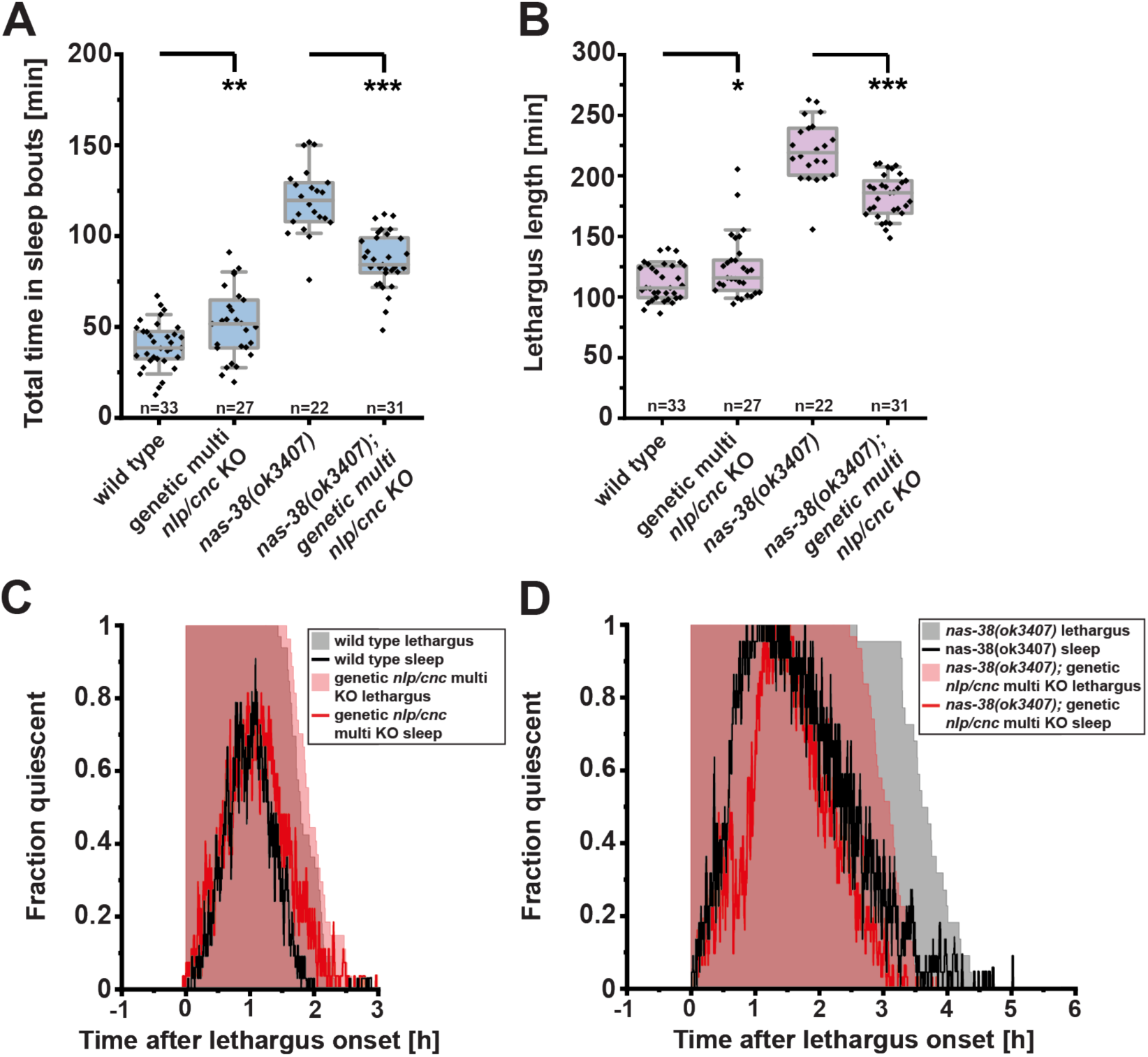
NAS-38 increases sleep through AMPs. (**A**) the *nlp* and *cnc* multi gene knockout suppressed increased sleep in *nas-38(ok3407)* by 27.6 %. (**B**) the *nlp* and *cnc* multi gene knockout suppressed prolonged lethargus duration in *nas-38(ok3407)* by 17.0 %. ***denotes p ≤ 0.001, ** denotes p ≤ 0.01, * denotes p ≤ 0.05, two-sample t-test. The boxplots show individual data points, the box represents the 25 % – 75 % range, the whiskers cover the 10 % - 90 % range, the thin gray line is the median. (**C**-**D**) fraction of worms in sleep during lethargus for the *nlp* and *cnc* multiple gene knockout experiment plotted as a function of time. The lines in intense colors indicate the sleep fraction, the pale shaded background indicates the fraction of worms in lethargus.

### Wounding signals through NLPs and CNCs to the nervous system to increase sleep

Many types of cellular stress, including heat, ethanol and wounding, increase sleep, a response that has been called stress-induced sleep and that is mediated by EGF signaling [15, 16, 54]. Increased sleep in the *nas-38(ok3407)* mutant is mediated by the epidermal innate immune response genes that are expressed after wounding. This suggests that the EGF signaling pathway is not the only pathway that signals damage, but that the innate immune pathway might increase sleep through NLP and CNC release following infection or wounding. We tested this hypothesis by wounding the cuticle and epidermis of adult worms with either a needle or a UV laser [28, 30]. We then placed the worms into microfluidic chambers and followed sleep behavior and RIS depolarization using fluorescence microscopy. Needle wounding as well as laser wounding triggered sleep bouts with a concomitant activation of RIS (Figure 6A-D for needle wounding, supplementary S6A-C for laser wounding). We next wanted to test whether *nas-38*-induced sleep during molting and wounding-induced sleep in the adult are regulated by the same molecular components. We hence measured sleep following mechanical wounding in *nas-38, sta-2*, and *nlp/cnc* mutants. *nas-38* mutations did not affect sleep, neither with nor without wounding (Figure S6D). *sta-2(ok1860)* reduced sleep after wounding by 58.7 % and the multi *nlp* and *cnc* deletion mutant suppressed wounding-induced sleep by 49.4 % (Figure 6D). We next tested whether wounding-induced sleep depended on RIS. For this, we wounded adult *aptf-1(gk794)* animals and quantified sleep. Wounding-induced sleep was abolished in RIS-deficient mutants (Figure 6D). These data indicate that *nas-38* gain-of-function during molting and mechanical injury of the adult epidermis present separate triggers that converge on innate immunity to cause sleep. Thus, we identified a novel type of sleep, innate immunity-induced sleep in *C. elegans*. Innate immunity-induced sleep requires the immune response (STA-2) and AMPs and is thus mechanistically distinct from EGF-induced sleep [15, 16, 54]. The partial suppression of wounding-induced sleep by immune response genes suggests that perhaps AMP and EGF signaling act in parallel to increase sleep following injury by converging on RIS neuron depolarization [55].

**Figure 6.**
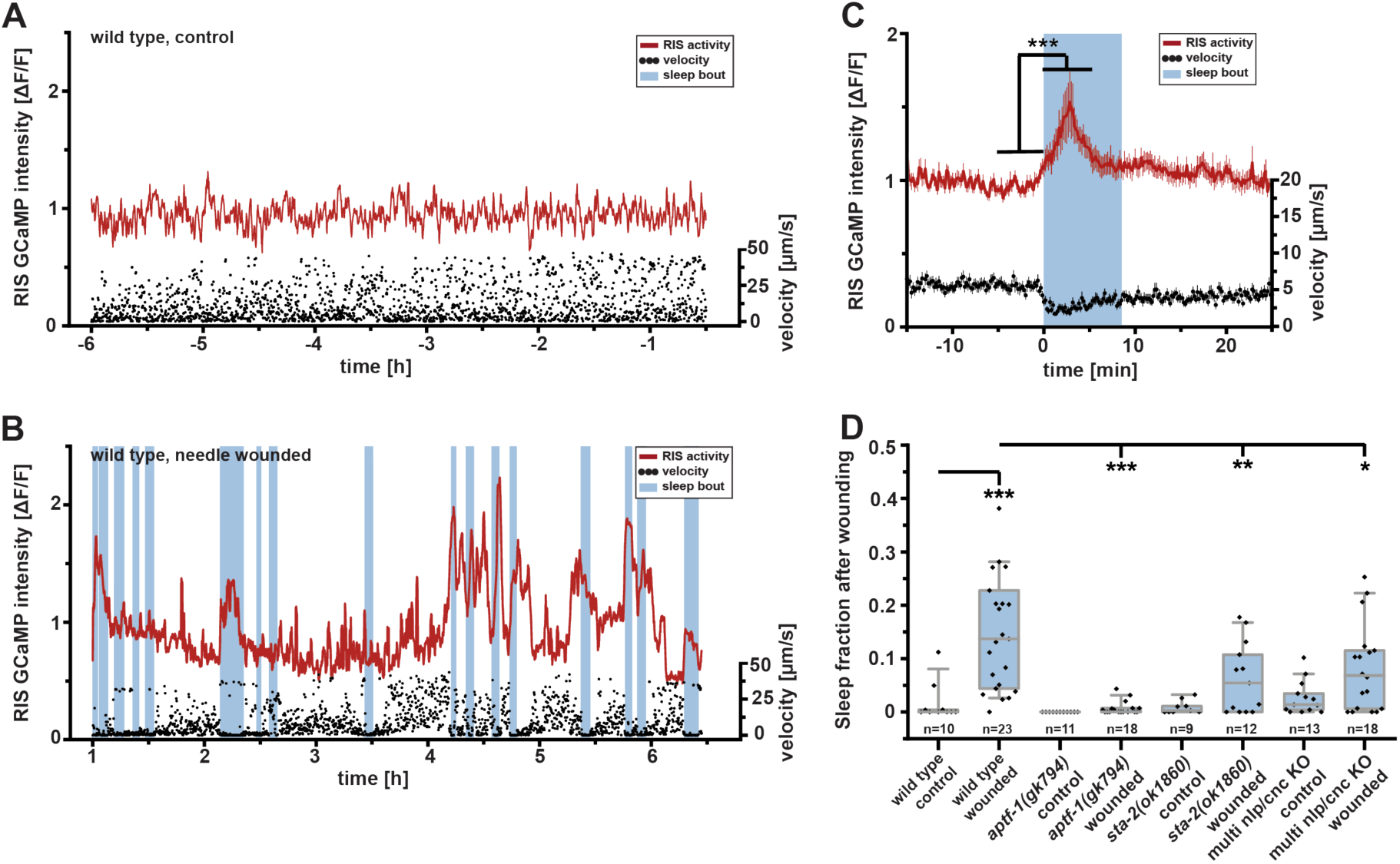
Wounding induces sleep via the innate immune response, AMPs, and the RIS neuron. (**A**) An individual trace of a wild type worm without wounding lacks strong RIS depolarization transients and sleep bouts. Representative plot showing velocity (black) and RIS calcium activity (red). Sleep bouts defined as locomotion quiescence are shown in blue. The 6h imaging time of control worms reflects the period in which sleep was scored immediately after wounding. (**B**) Data of an example wild type worm after sterile needle wounding. RIS activation transients and sleep bouts are visible. (**C**) Averaged velocity and RIS calcium activity after needle wounding aligned to sleep bout onset. RIS calcium activity (ΔF/F) increased by an average of 21.0 % during sleep bouts. n = 30 worms; *** p ≤ 0.001; paired Wilcoxon rank test. (**D**) Worms without wounding showed almost no sleep bouts. Needle-wounded wild type individuals slept 14.5 % of the time. *aptf-1(gk794)* reduced sleep by 95.2 %. *sta-2(ok1860)* suppressed sleep after wounding by 58.7 % and the *nlp* and *cnc* multi gene knockout suppressed sleep by 49.4 %. ***denotes p ≤ 0.001, ** denotes p ≤ 0.01, * denotes p ≤ 0.05, two-sample t-test. The boxplots show individual data points, the box represents the 25 % – 75 % range, the whiskers cover the 10 % - 90 % range, the thin gray line is the median.

### Many *NLPs* and *CNCs* can induce sleep through the RIS neuron

*nlp* and *cnc* genes are involved in wounding-induced sleep. This suggested that individual *nlp* and *cnc* peptides can induce sleep when expressed and released in adult worms. To test this hypothesis, we overexpressed *nlp* and *cnc* genes individually and measured sleep behavior. We conditionally and broadly overexpressed each of the *nlp* and *cnc* genes that were up regulated in *nas-38(ok3407)* by using a heat-shock promoter. To induce expression, adult worms were mildly heat shocked and the fraction of behaviorally quiescent animals was manually scored over time. For all 17 *nlp* and *cnc* genes tested, overexpression led to an induction of quiescence behavior after about 0.5-2.5 h that was not observed in control heat-shocked worms (Figure 7A and S7A-Q). To see whether NLPs and CNCs act through RIS, we tested quiescence induction by overexpression of *nlp* and *cnc* genes in *aptf-1(gk794). aptf-1(gk794)* efficiently suppressed most of the *nlp* and *cnc* overexpression-induced immobilization in sixteen out of seventeen peptides that were tested, and reduced it by almost half in *cnc-10* (Figure 7A and S7A-Q). Thus, the ability of *nlp* and *cnc* genes to induce sleep indicates that they function as somnogens. *NLPs* and *CNCs* act mostly through RIS to increase sleep.

**Figure 7.**
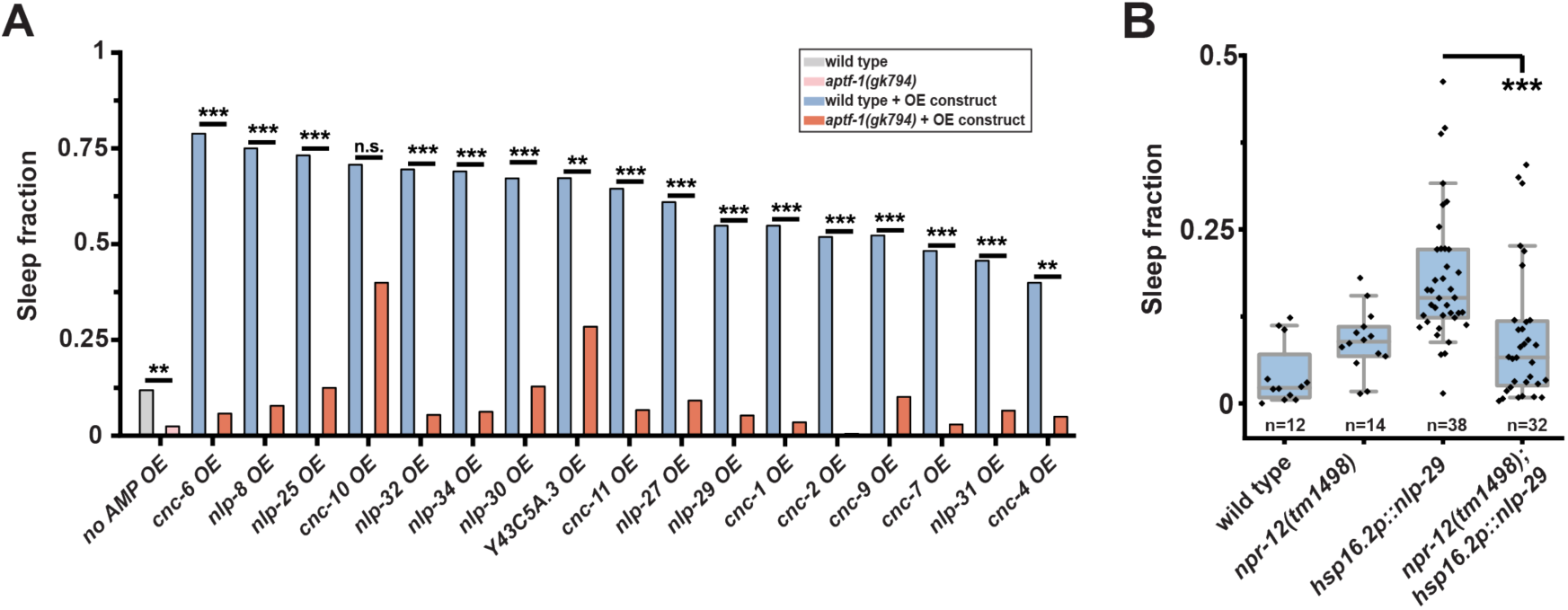
Many AMPs can induce sleep in the adult worm via RIS, and NLP-29-induced sleep requires the NPR-12 neuropeptide receptor. (**A**) Effect of *nlp* and *cnc* gene overexpression on behavioral quiescence in wild type and in *aptf-1(gk794)* background. Sleep was measured from 30 min till 150 min after the heat shock. *aptf-1(gk794)* reduced endogenous quiescence after heat shock by 79.4 %. *aptf-1(gk794)* also reduced AMP overexpression (OE)-induced sleep significantly in almost all conditions (by 92.7 % in *cnc-6* OE, 89.6 % in *nlp-8* OE, 82.9 % in *nlp-25* OE, 92.3 % in *nlp-32* OE, 91.0 % in *nlp-34* OE, 80.8 % in *nlp-30* OE, 57.7 % in Y43C5A.3 OE, 89.7 % in *cnc-11* OE, 84.9 % in *nlp-27* OE, 90.4 % in *nlp-29* OE, 93.6 % in *cnc-1* OE, 99.0 % in *cnc-2* OE, 80.6 % in *cnc-9* OE, 93.9 % in *cnc-7* OE, 80.9 % in *nlp-31* OE, and 87.5 % in *cnc-4* OE). *** denotes p ≤ 0.001, ** denotes p ≤ 0.01, Fisher’s exact test. (**B**) Knockout of the neuropeptide receptor gene *npr-12* suppresses *nlp-29* overexpression-induced sleep by 46.7 %. *** denotes p ≤ 0.001, two sample t-test. The boxplots show individual data points, the box represents the 25 % – 75 % range, the whiskers cover the 10 % - 90 % range, the thin gray line is the median.

### NLP-29 acts through the neuropeptide receptor NPR-12

How can AMPs signal to the nervous system to control RIS activity? NLP-29 has been shown to bind to and activate the NPR-12 neuropeptide receptor to control dendrite degeneration in sensory neurons during aging and infection [56]. We hence wanted to test whether NLP-29-induced sleep requires NPR-12. We crossed a loss-of-function mutation of NPR-12 into the NLP-29 overexpression strain, induced NLP-29 by a mild heat shock in adult worms and tested for suppression of sleep. In *npr-12(tm1498)* mutant animals, sleep amount was reduced by 46.7 % in the 0.5 – 3.25 h following the induction of NLP-29 overexpression (Figure 7B). Thus, NLP-29 increases sleep though NPR-12. The neuropeptide receptor is expressed in FLP and PVD sensory neurons as well as in PVC interneurons that control locomotion but not in RIS [56]. FLP and PVC neurons synapse onto RIS [57], and, intriguingly, the PVC neurons have recently been shown to be key activators of RIS [58]. Single-cell sequencing showed that NPR-12 is expressed also in RIM neurons [59], which are also involved in RIS activation [58]. However, NPR-12 is not expressed in RIS itself [55]. Thus, NLP-29 signals need for sleep to the nervous system through the NPR-12 neuropeptide receptor that is expressed in the brain. NLP-29 does not appear to activate RIS directly, but acts through NPR-12 in circuits that regulate RIS activity.

### Sleep increases the chance to survive injury

The increase of sleep by wounding suggests that sleep induction by innate immunity might increase the chance to survive injury. To test this idea, we wounded animals with a glass needle and tested whether individuals were still alive 24h and 48h after the insult. Wild-type animals showed a moderate percentage of dead individuals upon wounding (12% after 24h, 14% after 48h). By contrast, *aptf-1(gk794)*, in which sleep was impaired, had an almost tripled risk of dying following the injury (34% after 24h, p ≤ 0.01, 42% after 48h, p ≤ 0.001, Fishers’ exact test.) (Figure 8A-B). In addition, knockdown of *nlp/cnc* genes also reduced the chance to survive the injury. After 48h, worms lacking 19 *nlp* and *cnc* genes died significantly more often (32%, p ≤ 0.01, Fishers’ Exact test, Figure 8A-B). These data indicate that *nlp/cnc* genes and sleep provide an advantage for surviving injury, thus demonstrating a biological significance for innate immunity-induced sleep (Figure 8C).

**Figure 8.**
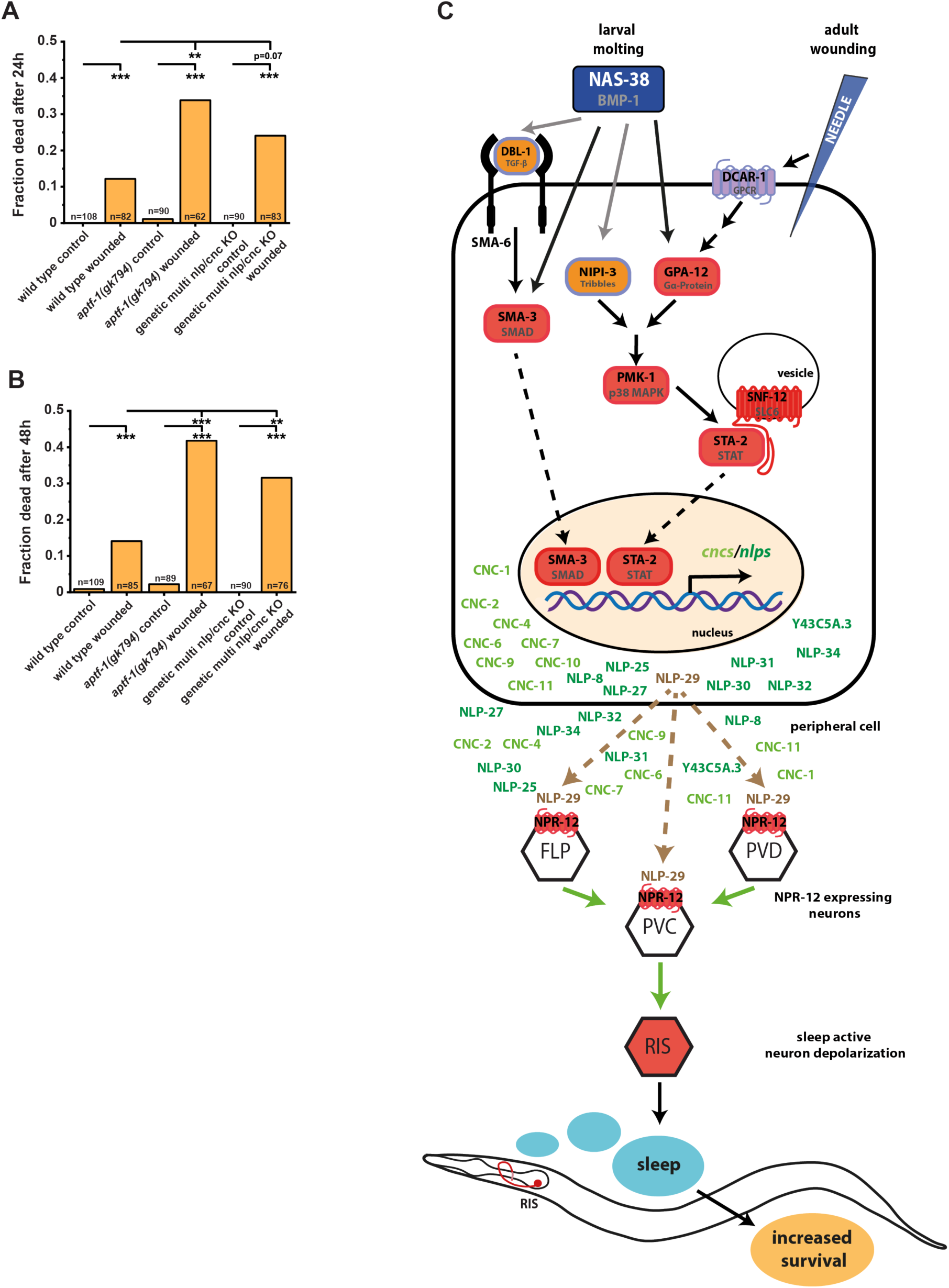
Sleep increases survival after wounding. (**A**) – (**B**) Needle wounding causes moderate mortality in young adults. Mortality rate was counted 24h and 48h after the injury. Sleepless *aptf-1(gk794)* mutant worms and worms lacking 19 AMP genes are significantly more likely to die from the injury than wild types. ***denotes p ≤ 0.001, ** denotes p ≤ 0.01, Fisher’s exact test. (**C**) Cartoon model for how NAS-38 and sterile wounding induce sleep via an innate immune response pathway. Activation of innate immunity pathways by NAS-38 in peripheral tissues leads to an increase of AMP expression and secretion. AMPs then activate receptors including NPR-12 in neuronal circuits that control RIS activity. Activation of RIS ultimately leads to sleep induction. Proteins corresponding to mutants that suppressed the induction of sleep are marked in red. Proteins corresponding to mutants that suppressed the induction of NLP-29 expression are shown in orange. Proteins corresponding to mutants that did not suppress are shown in violet.

## Discussion

### An innate immune response signals increased need for sleep caused by injury from peripheral tissues through AMPs and a neuropeptide receptor to sleep-inducing circuits

Tolloids are metalloproteases that are involved in collagen processing required for formation of the extracellular matrix. They are well known for cleaving the c-propeptide of collagen required for bone and cartilage formation [46, 60]. The role of tolloid-related factors in matrix processing appears to be conserved in *C. elegans*. For example, HCH-1 cleaves protein components of the eggshell to facilitate hatching [61], DPY-31 is required for collagen processing during cuticle formation [62], and NAS-36 and NAS-37 are involved in shedding of the old cuticle (ecdysis) [63, 64]. NAS-36 and NAS-37, like NAS-38, contain a thrombospondin domain. Tolloids/BMP-1-related proteins also play a second role, namely in developmental processes. Tolloid in *Drosophila* is well known as a developmental factor required for dorso-ventral patterning. It acts by cleaving decapentaplegic, a TGF-β homolog, which in turn leads to transcriptional changes. Similarly, BMP1 in mammals also cleaves TGF-β to control cartilage and bone development [46]. Here we show that NAS-38 activation triggers an innate immune response during larval molting. NAS-38, similar to Tolloid/BMP-1, acts through TGF-β/DBL-1 pathway component SMA-3, demonstrating that the canonical signaling components of the tolloid pathway have been conserved in *C. elegans*. NAS-38 signals through a heteromeric G alpha subunit, GPA-12 onto the p38 MAP kinase pathway. Both pathways lead to the expression of *nlp* and *cnc* genes, many of which act as AMPs, dependent on STA-2 in the epidermis [35]. These innate immune pathways are normally triggered by injury and infection. Thus, NAS-38 activation might act by causing an injury, perhaps through its protease function in collagen remodeling during molting. In addition, NAS-38 could act directly through activation of TGF-β signaling. During normal larval molting, NAS-38 promotes AMP expression, but knocking down AMPs did not reduce sleep. NAS-38 gain-of-function mutation strongly triggered the immune response during larval molting thus revealing the roles of immune genes and peripheral AMPs in sleep. This suggests that there are redundant pathways that promote sleep during molting, and one of these pathways appears to be an immune response promoted by NAS-38.

Seminal work showed that cellular stress increases sleep through EGF signaling [15, 16, 54]. We show that wounding increases sleep through activation of innate immunity and peripheral AMPs and we hypothesize that they may also play this role following infection. Thus, EGF-induced sleep and AMP-induced sleep might function in parallel following injury. In adult animals, innate immunity is triggered by wounding and infection [28, 35, 37]. The activation of STAT through wounding is suggested to be conserved in humans. Wounding of human epidermal keratinocytes activates the STAT proteins STAT3 and STAT5B and leads to the productions of AMPs such as β-defensins and pro-inflammatory cytokines (IL-6 and IL-8)[65]. Pro-inflammatory cytokines such as IL-1β, TNF-α, and IL-6 induce sleep and are involved in the increased sleepiness caused by infection [1, 2, 66]. AMPs can bind to and interfere with microbial membranes but also act as modulators of the immune system. The AMP encoded by *nemuri* has been shown to be expressed and act in the brain to promote sleep, perhaps by acting like a sleep-promoting neurotransmitter [18]. Here we describe an additional, novel role for a large group of AMPs. We show that AMPs can act as long-range somnogen messenger molecules that signal increased need for sleep from peripheral tissues, such as the epidermis, to the brain to promote sleep. We show that many AMPs act redundantly to promote sleep, and potential additional sleep-promoting AMPs exist in *C. elegans*. AMPs might also act as long-range somnogens in other organisms.

AMPs have been proposed to be secreted from the epidermis and to activate the NPR-12 neuropeptide receptor in the PVD sensory neuron, to modulate aging pathways in this neuron [56]. The NPR-12 receptor is expressed in circuits that include neurons that are known to control depolarization of RIS, but not in RIS itself, thus suggesting a model in which RIS is depolarized by NPR-12-expressing circuits, which are controlled by NLP-29. Whether other AMPs also activate NPR-12 or whether additional receptors exist that respond to specific AMPs are interesting questions for future studies. The high number of AMPs that we found to be able to promote sleep is striking and may reflect the biological importance of innate immunity-induced sleep. Speculatively, it could increase the robustness of the sleep response, and circumvent potential subversion of immune signaling by pathogens. In summary, we have discovered a long-range signaling mechanism that connects peripheral wounding to sleep induction via AMPs that signal sleep need from the site of injury to the brain, causing sleep-neuron activation and thus sleep (Figure 8C). Increasing sleep following injury provides a selective advantage as it almost triples the chance of surviving. AMPs might also serve as a long-range signaling cue to promote sleep following wounding to increase the chance of recuperation in other organisms including mammals.

## Acknowledgments

We thank Ines Lewandrowski, Sinem Öztas, Juliane Haase, and Shizue Omi for help with some experiments. We thank Dong Yan, National BioResource Project Japan, and the *Caenorhabditis* Genetics Center supported by the National Institutes of Health Office of Research Infrastructure Programs (P40 OD010440) for strains. The CRISPR strains were generated by SunyBiotech and Knudra. We thank Hirofumi Toda for discussing the role of *nemuri*. This work was supported by the Max Planck Society (Max Planck Research Group “Sleep and Waking”), by an European Research Council Starting Grant (ID: 637860, SLEEPCONTROL), the French National Research Agency (ANR-16-CE15-0001-01), by the « Investissements d’Avenir " French Government program (ANR-16-CONV-0001) and from Excellence Initiative of Aix-Marseille University - A*MIDEX and institutional grants from CNRS, Aix Marseille University, National institute of Health and Medical Research (Inserm) to the CIML.

## Material and Methods

### *C. elegans* maintenance and strains

*C. elegans* was grown under standard conditions [67]. Worms were maintained at 20°C on nematode growth medium (NGM) plates seeded with *E. coli* OP50, unless otherwise mentioned. The *C. elegans* strains used in this project are listed in the appendix (supplementary table 2). If not otherwise mentioned, the strains were ordered at the CGC or created for this project. The following strains were used:

| Strain | Genotype | Figure | Source |
| --- | --- | --- | --- |
| N2 | wild type | 1A, 1C, 1D-E, 1G-H, 2A-C, 3B-C, 5A-C, 6D, 7A-B, 8A-B, S1A-B, S2, S3A-B, S5A-D, S6C-D, S7A-Q | CGC |
| HBR1114 | *nas-38(tm2655) X.* | 1D-E, 2A, S6D | National BioResource Project, Japan |
| COP1622 | *nas-38(knu568) X.* | 1D-E, 3B, S3C | Knudra Transgenics for this project |
| HRBR1010 | *nas-38(ok3407) X.* | 1B-E, 1G-H, 2A-C, 3C, 5A-B, 5D, S1A-B, S2, S3A-B, S5A-D, S6D | This project |
| COP1635 | *nas-38(knu579ok3407) X.* | 1D-E | Knudra Transgenics for this project |
| HBR1110 | *unc-119(ed3) III; goels257[nas-38p::d1mgfp::nas-38-3'-UTR, unc-119(+)].* | 1I-J | This project |
| HBR227 | *aptf-1(gk794) II.* | 1G-H, 5E, 6D, 7A, 8A-B, S7A-Q | [22] |
| HBR1777 | *goels384 [flp-11p::egl-1::SL2-mKate2-flp-11-3'utr, unc-119(+)].* | 1G-H | [68] |
| HBR2072 | *nas-38(ok3407) X; goels384[flp-11p::egl-1::SL2-mKate2-flp-11-3'utr; unc-119(+)].* | 1G-H | This project |
| HBR977 | *aptf-1(gk794) II; nas-38 (ok3407) X; goels118[aptf-1-5'utr::SL1-GCaMP3.35-SL2::mKate2-aptf-1-3'utr,unc-119(+)].* | 1G-H | This project |
| HBR1089 | *frIs7[nlp-29p::GFP+coel-12p::dsRed] IV.* | 2D, 2F, 4A-F, S2B, S4A-G | This project |
| HBR1090 | *frIs7[nlp-29p::GFP, coel-12p::DsRed] IV, nas-38(ok3407) X.* | 2D, 2F, 4A-F, S2B, S4A-G | This project |
| IG1631 | *frSi11[pNP152(sta-2p::STA-2::mkate2), unc-119(+), ttTi5605] II.* | 2E, 2G | This project |
| HBR1925 | *frSi11[pNP152(sta-2p::STA-2::mkate2); unc-119(+), ttTi5605] II; nas-38(ok3407) X.* | 2E, 2G | This project |
| NU3 | *dbl-1(nk3) V.* | 3B-C | CGC |
| CB491 | *sma-3(e491) III.* | 3B-C | CGC |
| KU25 | *pmk-1(km25) IV.* | 3B-C | CGC |
| IG1241 | *sta-2(ok1860) V.* | 3B-C | [35] |
| IG1242 | *frIs7[nlp-29p::GFP, coel-12p::DsRed] IV; snf-12(tm692) X.* | 3B-C, 4D-E | [35] |
| HBR2010 | *sma-3(e491) III; pmk-1(km25), frIs7[nlp-29p::GFP, coel-12p::DsRed] IV.* | S3A-B, S4F-G | This project |
| IG460 | *pmk-1(km25), frIs7[nlp-29p::GFP, coel-12p::DsRed] IV.* | S4E, S4G | [28] |
| IG342 | *frIs7[nlp-29p::GFP, coel-12p::DsRed] IV; nipi-3(fr4) X.* | S4D, S4G | [28] |
| IG722 | *frIs7[nlp-29p::GFP, coel-12p::DsRed] IV; gpa-12(pk322) X.* | 4B, 4E, S3A-B | [36] |
| IG1424 | *frIs7(nlp-29p::GFP, coel-12p::DsRed) IV; dcar-1(tm2484) V.* | S3A-B, S4C, S4G | [29] |
| HBR2146 | *sma-3(e491) III; frIs7[nlp-29p::GFP, coel-12p::DsRed] IV.* | S4B, S4G | This project |
| HBR2127 | *frIs7[nlp-29p::GFP, coel-12p::DsRed] IV; dbl-1(nk3)V.* | S4A, S4G | This project |
| HBR2375 | *frIs7[nlp-29p::GFP, coel-12p::DsRed] IV; sta-2(ok1860)V.* | 4C, 4F | This project |
| HBR2186 | *frIs7[nlp-29p::GFP, coel-12p::DsRed] IV; dbl-1(nk3) V; nas-38(knu568) X.* | 3B, S3C | This project |
| HBR1923 | *frIs7[nlp-29p::GFP, coel-12p::DsRed] IV; dbl-1(nk3) V; nas-38(ok3407) X.* | 3B-C, 4F,, S4A, S4G | This project |
| HBR1884 | *sma-3(e491) III; frIs7[nlp-29p::GFP, coel-12p::dsRed] IV; nas-38(ok3407) X.* | 3B-C, 4F S4B, S4G | This project |
| IG1670 | *frIs7[nlp-29p::GFP, coel-12p::DsRed] IV; dcar-1(tm2484) V; nas-38(ok3407) X.* | 4F, S3A-B, S4C, S4G | This project |
| HBR1877 | *frIs7[nlp-29p::GFP, coel-12p::DsRed] IV; gpa-12(pk322), nas-38(ok3407) X.* | 4B, 4E-F, S3A-B | This project |
| IG1676 | *frIs7[nlp-29p::GFP, coel-12p::DsRed] IV; nas-38(ok3407), nipi-3(fr4) X.* | 4F, S3A-B, S4D, S4G | This project |
| HBR1878 | *pmk-1(km25) IV; nas-38(ok3407) X.* | 3B-C, 4F | This project |
| HBR2376 | *pmk-1(km25), frIs7 [nlp-29p::GFP, coel-12p::DsRed] IV; nas-38(ok3407) X.* | S4E, S4G | This project |
| HBR1541 | *sta-2(ok1860) V; nas-38(ok3407) X.* | 3A-B, 4F | This project |
| IG1803 | *frIs7[nlp-29p::GFP, coel-12p::DsRed] IV; nas-38(ok3407), snf-12(tm692) X.* | 3B-C, 4D-F, | This project |
| HBR2014 | *sma-3(e491) III; pmk-1(km25), frIs7[nlp-29p::GFP, coel-12p::DsRed] IV; nas-38(ok3407) X.* | 4F, S3A-B, S4F-G | This project |
| HBR2145 | *frIs7[nlp-29p::GFP, coel-12p::DsRed] IV; nas-38(knu568) X.* | 4A, 4F | This project |
| HBR1117 | *frIs7[nlp-29p::GFP + coel-12p::DsRed] IV; nas-38 (tm2655) X.* | 4A | This project |
| HBR2185 | *sma-3(e491) III; frIs7[nlp-29p::GFP, coel-12p::DsRed]IV; nas-38(knu568) X.* | 3B, 4F, S3C | This project |
| HBR1928 | *frIs7[nlp-29p::GFP, coel-12p::DsRed] IV; sta-2(ok1860) V; nas-38(ok3407) X.* | 4C, 4E-F | This project |
| HBR2196 | *pmk-1(km25) IV; nas-38(knu568) X.* | 3B, 4F, S3C | This project |
| HBR2090 | *sta-2(ok1860) V; nas-38(knu568) X.* | 3B, 4F, S3C | This project |
| HBR2190 | *frIs7[nlp-29p::GFP, coel-12p::DsRed] IV; nas-38(knu568), snf-12(tm692) X.* | 3B, 4F, S3C | This project |
| PHX293 | *nas-38(syb293) X.* | S1A-B | SunyBiotech for this project |
| HBR1272 | *nlp-29 (tm1931) V; nas-38(ok3407) X.* | S5A-B | This project |
| HBR1556 | *nlp-29(tm1931) V.* | S5A-B | This project |
| HBR2317 | *nlp-8(syb762) IV.* | S5A-B | This project |
| HBR2318 | *nlp-8(syb762) IV; nas-38(ok3407) X.* | S5A-B | This project |
| PHX1446 | *nlp-8(syb762) I; nlp-32(syb431), cnc-6(syb393) III, Y43C5A.3(syb761) IV; sybDf1, sybDf2, cnc-7(syb558), cnc-10(syb406), nlp-25(syb579) V.* | 5A-C | SunyBiotech for this project |
| PHX1521 | *nlp-8(syb762) I; nlp-32(syb431), cnc-6(syb393) III, Y43C5A.3(syb761) IV; sybDf1, sybDf2, cnc-7(syb558), cnc-10(syb406), nlp-25(syb579) V; nas-38(ok3407) X.* | 5A-B, 5D | SunyBiotech for this project |
| HBR2301 | *npr-12(tm1498) IV.* | 7B | *tm1498* provided by Dong Yan |
| HBR2302 | *npr-12(tm1498) IV; goels326[hsp-16.2p::nlp-29::SL2-mkate2-unc-54-3'UTR, unc-119(+)]*. | 7B | This Project |
| HBR1892 | *aptf-1(gk794) II; goels388[hsp-16.2p::cnc-1::SL2-mkate2-unc-54-3'utr, unc-119(+)]*. | 7A, S7A | This project |
| HBR1896 | *goels388[hsp-16.2p::cnc-1::SL2-mkate2-unc-54-3'utr, unc-119(+)]*. | 7A, S7A | This project |
| HBR1893 | *aptf-1(gk794) II; goels397[hsp-16.2p::cnc-10::SL2-mkate2-unc-54-3'utr, unc-119(+)]*. | 7A, S7G | This project |
| HBR1897 | *goels397[hsp-16.2p::cnc-10::SL2-mkate2-unc-54-3'utr, unc-119(+)]*. | 7A, S7G | This project |
| HBR1905 | *goeEx672[hsp-16.2p::cnc-11::SL2-mkate2-unc-54-3'utr, unc-119(+)]*. | 7A, S7H | This project |
| HBR1906 | *aptf-1(gk794) II; goeEx672[hsp-16.2p::cnc-11::SL2-mkate2-unc-54-3'utr, unc-119(+)]*. | 7A, S7H | This project |
| HBR1895 | *aptf-1(gk794) II; goels407[hsp-16.2p::cnc-2::SL2-mkate2-unc-54-3'utr, unc-119(+)]*. | 7A, S7B | This project |
| HBR1901 | *goels407[hsp-16.2p::cnc-2::SL2-mkate2-unc-54-3'utr, unc-119(+)]*. | 7A, S7B | This project |
| HBR1911 | *goels411[hsp-16.2p::cnc-4::SL2-mkate2-unc-54-3'utr, unc-119(+)]*. | 7A, S7C | This project |
| HBR1960 | *aptf-1(gk794) II; goels411[hsp-16.2p::cnc-4::SL2-mkate2-unc-54-3'utr, unc-119(+)]*. | 7A, S7C | This project |
| HBR1907 | *goels410[hsp-16.2p::cnc-6::SL2-mkate2-unc-54-3'utr, unc-119(+)]*. | 7A, S7D | This project |
| HBR1908 | *aptf-1(gk794) II; goels410[hsp-16.2p::cnc-6::SL2-mkate2-unc-54-3'utr, unc-119(+)]*. | 7A, S7D | This project |
| HBR1912 | *goels412[hsp-16.2p::cnc-7::SL2-mkate2-unc-54-3'utr, unc-119(+)]*. | 7A, S7E | This project |
| HBR1913 | *aptf-1(gk794) II; goels412[hsp-16.2p::cnc-7::SL2-mkate2-unc-54-3'utr, unc-119(+)]*. | 7A, S7E | This project |
| HBR1914 | *goels413[hsp-16.2p::cnc-9::SL2-mkate2-unc-54-3'utr, unc-119(+)]*. | 7A, S7F | This project |
| HBR1915 | *aptf-1(gk794) II; goels413[hsp-16.2p::cnc-9::SL2-mkate2-unc-54-3'utr, unc-119(+)]*. | 7A, S7F | This project |
| HBR1963 | *goeEx657[hsp-16.2p::Y43C5A.3::SL2-mkate2-unc-54-3'utr, unc-119(+)]*. | 7A, S7I | This project |
| HBR1964 | *aptf-1(gk794) II; goeEx657[hsp-16.2p::Y43C5A.3::SL2-mkate2-unc-54-3'utr, unc-119(+)]*. | 7A, S7I | This project |
| HBR1903 | *goeEx667[hsp-16.2p::nlp-8::SL2-mkate2-unc-54-3'utr, unc-119(+)]*. | 7A, S7J | This project |
| HBR1904 | *aptf-1(gk794) II; goeEx667[hsp-16.2p::nlp-8::SL2-mkate2-unc-54-3'utr, unc-119(+)]*. | 7A, S7J | This project |
| HBR1961 | *goels431[hsp-16.2p::nlp-25::SL2-mkate2-unc-54-3'utr, unc-119(+)]*. | 7A, S7K | This project |
| HBR1962 | *aptf-1(gk794) II; goels431[hsp-16.2p::nlp-25::SL2-mkate2-unc-54-3'utr, unc-119(+)]*. | 7A, S7K | This project |
| HBR1900 | *goels408[hsp-16.2p::nlp-27::SL2-mkate2-unc-54-3'utr, unc-119(+)]*. | 7A, S7L | This project |
| HBR1957 | *aptf-1(gk794) II; goels408[hsp-16.2p::nlp-27::SL2-mkate2-unc-54-3'utr, unc-119(+)]*. | 7A, S7L | This project |
| HBR1549 | *goels326[hsp-16.2p::nlp-29::SL2-mkate2-unc-54-3'UTR, unc-119(+)]*. | 7A-B, S7M | This project |
| HBR1547 | *aptf-1(gk794) II; goels326[hsp-16.2p::nlp-29::SL2-mkate2-unc-54-3'UTR, unc-119(+)]*. | 7A, S7M | This project |
| HBR1909 | *goeEx676[hsp-16.2p::nlp-30::SL2-mkate2-unc-54-3'utr, unc-119(+)]*. | 7A, S7N | This project |
| HBR1910 | *aptf-1(gk794) II; goeEx676[hsp-16.2p::nlp-30::SL2-mkate2-unc-54-3'utr, unc-119(+)]*. | 7A, S7N | This project |
| HBR1894 | *aptf-1(gk794) II; goels406[hsp-16.2p::nlp-31::SL2-mkate2-unc-54-3'utr, unc-119(+)]*. | 7A, S7O | This project |
| HBR1899 | *goels406[hsp-16.2p::nlp-31::SL2-mkate2-unc-54-3'utr, unc-119(+)]*. | 7A, S7O | This project |
| HBR1902 | *goels409[hsp-16.2p::nlp-32::SL2-mkate2-unc-54-3'utr, unc-119(+)]*. | 7A, S7P | This project |
| HBR1958 | *aptf-1(gk794) II; goels409[hsp-16.2p::nlp-32::SL2-mkate2-unc-54-3'utr, unc-119(+)]*. | 7A, S7P | This project |
| HBR1959 | *goeEx651[hsp-16.2p::nlp-34::SL2-mkate2-unc-54-3'utr, unc-119(+)]*; *aptf-1(gk794) II*. | 7A, S7Q | This project |
| HBR1898 | *goeEx651[hsp-16.2p::nlp-34::SL2-mkate2-unc-54-3'utr, unc-119(+)]*. | 7A, S7Q | This project |
| HBR1361 | *goels304[flp-11p::SL1-GCaMP3.35-SL2::mKate2-unc-54-3'UTR, unc-119(+)]*. | 6A-C, S6A-C | [68] |

### Creating transgenic animals

By using the Three Fragment Gateway System (Invitrogen) [69], constructs were cloned into the pCG150 Vector that contains an *unc-119(+)* rescue sequence. To verify the correct sequence of the cloned constructs, plasmids were Sanger-sequenced [70]. Transgenes were expression-optimized for *C. elegans* [71]. We generated the transgenic strains by micro particle bombardment into *unc-119(ed3)* mutant worms and used a phenotypical rescue of the uncoordinated phenotype as a selection marker [72, 73]. The insertions that were obtained were backcrossed two times against N2 to remove the *unc-119(ed3)* background. The constructs created for this project are listed in the appendix (supplementary table 3).

*frSi11* is a single copy insertion on chromosome II at the location of the Nemagenetag Mos1 insertion [74] ttTi5605 of pNP152 (*col-62p::Lifeact::mKate2-c-nmy3’utr*). pNP152 was obtained by insertion of the *sta-2* promoter and *sta-2* gene fused to *Lifeact::mKate2* [75] into the MosSCI vector pCFJ151 [76] using Gibson Assembly (NEB Inc., MA) and confirmed by PCR or sequencing. It was injected into the EG6699 strain at 20 ng/µl together with pCFJ90 (myo-2p::mCherry) at 1.25 ng/µl, pCFJ104 (myo-3p::mCherry) at 5 ng/µl, pMA122 (hsp16.41p::PEEL-1) at 10 ng/µl, pCFJ601 (eft-3p::Mos1 transposase) at 20 ng/µl and pNP21 (*unc-53pB::GFP* [77]) at 40 ng/µl. A strain containing the insertion was obtained following standard selection and PCR confirmation [76]. The strain was then outcrossed with N2 male to remove the *unc-119* mutation.

|  |  |
| --- | --- |
| K185 | *nas-38p::d1mgfp::nas-38-3'-utr* |
| K281 | *flp-11p::egl-1::SL2-mkate2-flp-11-3'utr* |
| K82 | *aptf-1-5'utr::SL1-GCaMP3.35-SL2::mKate2-aptf-1-3'utr* |
| pNP152 | *sta-2p::sta-2::mkate2* |
| K271 | *hsp-16.2p::nlp-29::SL2-mkate2-unc-54-3'utr* |
| K305 | *hsp-16.2p::cnc-1::SL2-mkate2-unc-54-3'utr* |
| K304 | *hsp-16.2p::cnc-10::SL2-mkate2-unc-54-3'utr* |
| K284 | *hsp-16.2p::cnc-11::SL2-mkate2-unc-54-3'utr* |
| K282 | *hsp-16.2p::cnc-2::SL2-mkate2-unc-54-3'utr* |
| K294 | *hsp-16.2p::cnc-4::SL2-mkate2-unc-54-3'utr* |
| K283 | *hsp-16.2p::cnc-6::SL2-mkate2-unc-54-3'utr* |
| K295 | *hsp-16.2p::cnc-7::SL2-mkate2-unc-54-3'utr* |
| K296 | *hsp-16.2p::cnc-9::SL2-mkate2-unc-54-3'utr* |
| K292 | *hsp-16.2p::Y43C5A.3::SL2-mkate2-unc-54-3'utr* |
| K285 | *hsp-16.2p::nlp-8::SL2-mkate2-unc-54-3'utr* |
| K286 | *hsp-16.2p::nlp-25::SL2-mkate2-unc-54-3'utr* |
| K287 | *hsp-16.2p::nlp-27::SL2-mkate2-unc-54-3'utr* |
| K288 | *hsp-16.2p::nlp-30::SL2-mkate2-unc-54-3'utr* |
| K289 | *hsp-16.2p::nlp-31::SL2-mkate2-unc-54-3'utr* |
| K290 | *hsp-16.2p::nlp-32::SL2-mkate2-unc-54-3'utr* |
| K291 | *hsp-16.2p::nlp-34::SL2-mkate2-unc-54-3'utr* |
| K216 | *flp-11p::SL1-GCaMP3.35-SL2::mKate2-unc-54-3'utr* |

### Crossing *C. elegans* strains

*C. elegans* strains that carry multiple mutations or transgenes were created by conventional crossings and genotyped phenotypically or by a three-primer polymerase chain reaction (PCR) approach. The primer pairs used in this project are listed below (supplementary table 4):

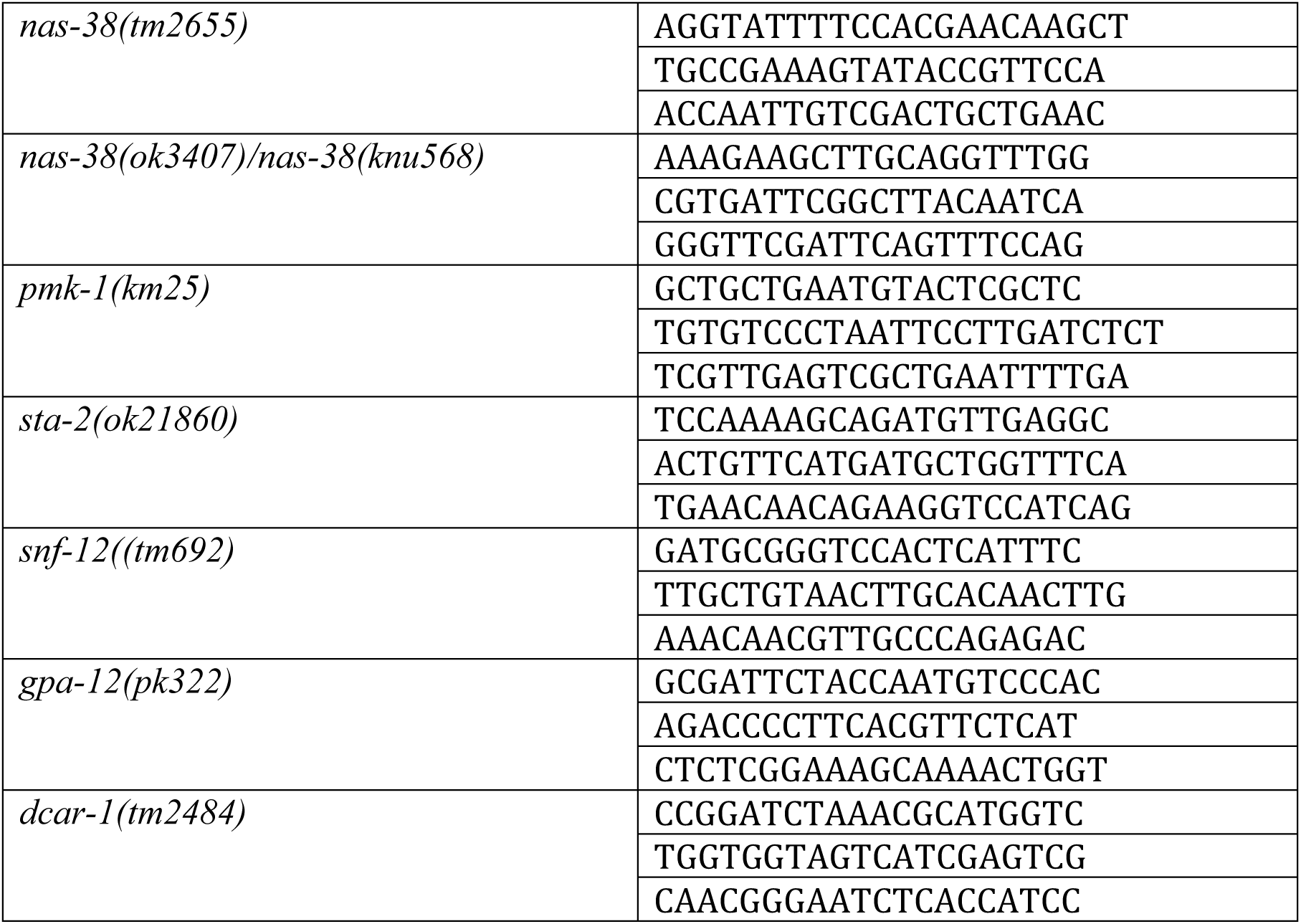
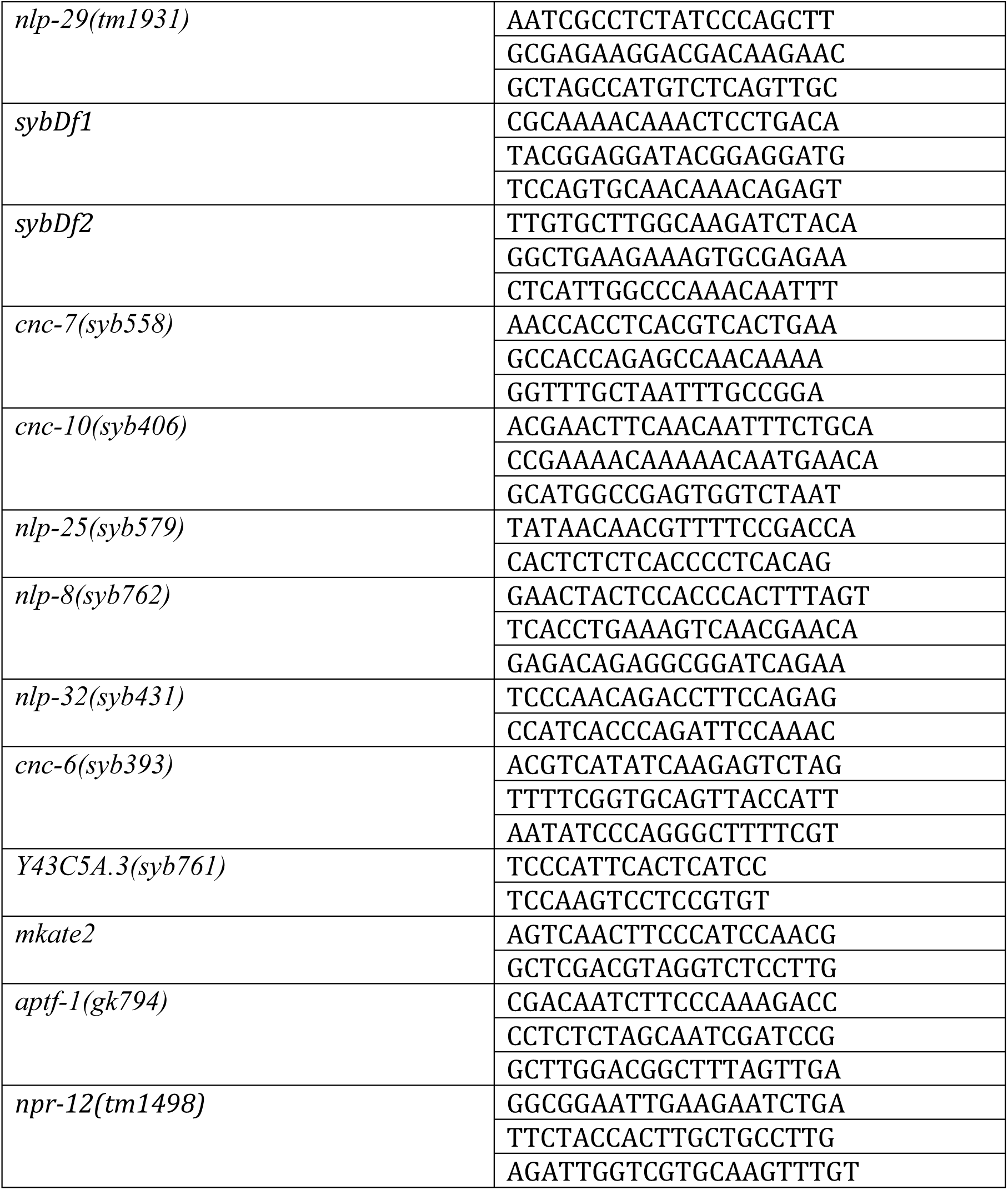

### Creating mutants with the CRISPR/Cas9 system

Strain generation with mutations in the endogenous locus via CRISPR/Cas9 was performed by Knudra Transgenics and SunyBiotech. In the multi *nlp* and *cnc gene* knockout strain 16 out of 17 *nlp*s and *cnc*s genes were deleted that were significantly upregulated in the *nas-38(ok3407)* mutant. As four of the significantly upregulated *cnc* genes and five of the *nlp* genes are organized in gene cassettes, we decided to delete those two cassettes completely to facilitate the generation of the multi knockout strain. This resulted in the knockout of three additional cassette members *cnc-3, cnc-5* and *nlp-28* that were all found upregulated in the *nas-38(ok3407)* background without reaching statistical significance. As both cassettes and 4 additional AMPs, namely *nlp-25, cnc-7, cnc-9* and *cnc-10*, are located on chromosome V and because the 5 deletions on this chromosome were already difficult to cross due to frequent recombination, we decided to forego the knockout of *cnc-9. cnc-9* was also not found to be upregulated upon infection [78].

### Genetic screen for mutants with increased behavioral quiescence

We obtained a mutant collection from a strain depository consisting of about 4500 strains with each strain harboring at least one known severe mutation such as a deletion or mutation associated with a phenotype. Because each strain had a mutation in a different gene, the mutant set covered strong alleles for at least 4500 different genes. Individual strains were grown on Nematode Growth Medium plates to obtain a mixed population of about 200-300 individuals containing all developmental stages including animals that were in lethargus. To identify candidate mutants that have increased sleep behavior, the plates where screened visually using a stereomicroscope for the presence of increased fraction of immobile larvae within the population (Supplementary Table 1).

### Imaging of *C. elegans* behavior

All long-term behavioral and functional Ca^2+^-imaging experiments presented in this project were done in agarose microchambers [23, 44]. The microchambers were created from 3-5 % hot melting agarose (Fisher-Scientific) dissolved in S-Basal or M9. The hot agarose was then cast into a polydimethylsiloxan (PDMS) mold to create micro-compartments in the agarose. After solidification of the agarose, the PDMS mold was removed and the chambers were filled with one worm (eggs or adults) each.

Needle-wounded adult worms were filmed in 700 µm x 700 µm x 45 µm or 370 µm x 370 µm x 45 µm (X length x Y length x Z depth) microchambers. Laser-wounded adult worms were filmed in 370 µm x 370 µm x 45 µm microchambers. L1 larvae were imaged in 190 µm x 190 µm x 15 µm microchambers. For 700 µm x 700 µm x 45 µm microchambers, the 10x objective was used. Adults in the 370 µm x 370 µm x 45 µm chamber and L1s were imaged with a 10x or 20x objective. L1 lethargus in the differential interference contrast (DIC) burst mode (explained below) was imaged with a 40x objective.

The microscope setup consisted of the following components: Either a Nikon TiE inverted microscope was used that was equipped with an automated XY stage (Prior, Nikon), an LED system (CoolLED), Andor iXon electron multiplying charge-coupled device (EMCCD) camera (512×512 pixels), kinetic trigger device, automated Prior XY stage, Andor iQ2 or iQ3 software and a self-made sample holder with a heat lid. Or a Nikon Ti2 inverted microscope was used that was equipped with an LED system (Lumencor Sola II), a Photometrics Prime 95B sCMOS camera (1608×1608 pixels), NIS Elements Advanced Research software and a custom-made sample holder with a heating lid. For fluorescence imaging, the transistor-transistor logic (TTL) trigger “fire out” of the camera was used to trigger the LED illumination. This decreased illumination time, because the sample was only illuminated during image acquisition.

### DIC imaging

L1 lethargus was measured in DIC burst or continues mode. In burst mode, every 15 or 30 min an acquisition burst with either 20 or 40 frames with a frame rate of 2 frames/s was recorded. Up to 60 worms were filmed in parallel. The DIC images were used to determine lethargus onset and the lethargus length based on the continuous absence of pharyngeal pumping. For continuous imaging, a frame rate of 0.2 frames/s was used for imaging L1 lethargus. Typically, 3-5 individual fields were filmed in parallel and up to 4 worms could be filmed in one field using the Andor EMCCD Camera and a 20x objective. The larger camera chip of the Photometrics Prime 95B allowed imaging 16 worms in one field using the 10x objective. Lethargus onset and end were determined optically by cessation and restart of feeding behavior. For adult wounding or heat shock experiments, a frame rate of 0.1 frames/s was used and only 1 worm was imaged per field.

### Functional Ca^2+^imaging

Standard GFP filter sets were used, the exposure time was set to 5 ms and gain to 100. 490 nm light intensity was 2.00 mW/mm^2^ using a 20x objective and 0.60 mW/mm^2^ using a 10x objective. The frame rate used was 0.1 frame/s.

### Fluorescent imaging experiments

We used fluorescent imaging in combination with the DIC burst mode to correlate gene expression with L1 lethargus onset, identified by cessation of pumping. Every 30 min, a DIC burst was recorded followed by a single fluorescent image. The *nlp-29p::GFP* transcriptional reporter was recorded with standard GFP filter sets in L1 larvae, the exposure time was set to 5 ms and gain was set to 30. A single image was taken using 490nm light with an intensity of 0.41 mW/mm^2^ using a 40x objective. If necessary, we used a neutral-density filter (ND4) to reduce excitation light intensity. For analysis of signal intensity, we used a lower intensity threshold of 250 arbitrary intensity units. In the L4 larvae, *nlp-29p::GFP* was imaged at the Ti2 setup using a 20x objective and an additional 1.5 x internal lens. The exposure time was set to 1ms and the light intensity was set to 17.32 mW/mm^2^.

The *nas-38p::d1GFP* transcriptional reporter was recorded with standard GFP filter sets, the exposure time was set to 5 ms and gain to 200. A z-stack was recorded with a plane distance of 1µm covering an overall distance of 20 µm. The 490 nm light intensity was 2.44 mW/mm^2^ using a 40x objective. After acquisition, the maximum projection of the z-stack was calculated. For analysis the lower threshold was set to 3000 intensity units. The *sta-2p::sta-2::mkate2* translational reporter was recorded using standard TexasRed filter sets in the L4 larvae using a spinning disk setup and a 100x objective. The exposure time was set to 100 ms and the gain was set to 300.

### Image analysis

#### Analyzing functional Ca^2+^ images

From functional Ca^2+^ sensor images, we extracted two parameters: 1) the worm’s locomotion velocity based on the movement of the neuron center and 2) the neuronal activity. Both parameters were determined by a custom-made MatLab (MathWorks) routine, similar to the one previously described [23].

For RIS depolarizing, the RIS activity was the average of the 30 highest pixel intensities in an 11 pixels x 11 pixel area around the detected pixel with the highest intensity. These 30 pixels represent the RIS cell body very robustly, which was determined empirically.

#### Analysing *sta-2::mKate2* localization in nuclei

Fluorescent signals of the *sta-2::mKate2* reporter in nuclei was identified using the inbuilt circle finding function *imfindcircles* in MatLab (MathWorks) with the nuclei radius range set from 11 – 30 pixels and a sensitivity of 0.92.

#### Fluorescent reporter analyses

Analysis of *nlp-29p::GFP* expression in Figure 2F was quantified with the COPAS Biosort (Union Biometrica; Holliston, MA) as described in [39]. The ratio between GFP intensity and size (time of flight; TOF) is represented in arbitrary units.

#### Sleep bout detection

For wounded adults under fed conditions, a quiescence bout had to last for at least 3 min to be classified as a sleep bout. Quiescence bouts during L1 lethargus and in starved wounding conditions had to have a minimum duration of 1 min to be classified as a sleep bout and quiescence bouts of adults following *nlp-29* overexpression had to last for at least 30 s to be classified as a sleep bout. Empirically determined cut-off parameters were used for sleep bout detection from image subtraction data. Frame subtraction values were smoothed using a 1^st^ degree polynomial local regression model over 5 time points and velocity data (only used in tracking RIS GCaMP after wounding experiments) was smoothed over 25 data points. This was done using the in-build *smooth* function in MatLab. The smoothed velocity values had to be below 0.5 µm/s for adults. When smoothed and normalized image subtraction data was used, the threshold for sleep bouts was set to 0.2 for L1 lethargus and starved wounded adults and to 0.5 for adults after heat shock induced *nlp-29* overexpression. As *nas-38(ok3407)* mutants recover more slowly from lethargus, phases of decreased mobility can be detected in rare cases also outside of lethargus. These are not counted as sleep bouts for further analysis.

#### L1 and lethargus duration

The duration of the entire L1 stage and the lethargus period were measured in microchambers. Using a DIC burst protocol, the following time points were manually extracted: the onset of hatching, the onset and end of lethargus (defined by the cessation and resumption of pharyngeal pumping) and the shedding of the cuticle. From these time points the duration of the L1 stage was determined as the time between hatching and L1 cuticle shedding and lethargus length was determined as the period before cuticle shedding during which animals did not feed.

#### Heat shock

To induce overexpression of AMPs, we generated transgenes that drive expression of the *cnc* or *nlp* gene from the *hsp-16.2* promotor. Expression from this promoter is strongly increased at temperatures above 33°C [79]. For the screen of sleep induction by overexpression of the individual AMPs, worms were picked to small NGM plates and their sleep-behavior quantified. The plates were sealed tightly using Parafilm (Pechiney Plastic Packaging). After 10 min of heat shock in a water bath at 37°C the plates were placed on crushed ice for 2 min, to stop the heat shock. When removed from the ice, this time point was defined as 0 h and the number of sleeping worms was scored manually. Scoring was repeated in 30 min intervals for at least two hours as described [19, 25].

For testing for suppression of the *nlp-29* OE-induced sleep with the NPR-12 receptor gene knockout, a home-made temperature control device with a Peltier element was used as described before [55]. In brief, up to 15 young adult worms were transferred into agarose chambers (370 µm x 370 µm x 45 µm) without food. We sealed the microchambers with a coverslip, thereby distributing the worms into individual microchambers. The coverslip was then attached with sticky tape to a metal plate of the temperature control device that had an opening cut-out, and the device was mounted onto the microscope. For applying the heat shock, the heater was set to 38 °C, resulting in 37 °C in the chamber for 15 min, as measured by a thermoelement. 3 min before the heat shock, the heating lid was set to 39.5 °C. Before and after the heat shock period, the heating lid was set to 25.5 °C and the heater was turned off, resulting in room temperature of the sample inside the agarose chamber. During the experiment, each worm was imaged every 10 s and for at least 4 h. For quantifying quiescence behavior, the first 30 min before the heat shock were used as baseline and 4 h after heat shock were used to determine sleep induced by the overexpression.

#### *C. elegans* RNAi-by-feeding

RNAi expressing plasmids used were based on the LH4440 feeding vector [80]. The plasmids were transformed into the HT115 *E. coli* strain and RNA expression was induced by 1 mM isopropyl-β-d-thiogalactopyranoside (IPTG, Sigma-Aldrich). We used a slightly modified version of the established RNAi-by-feeding protocol [81]. *E. coli* were cultured overnight in LB medium (50 µg/ml ampicillin, Roth) and then seeded to freshly made NMG plates containing 1mM of IPTG (Sigma-Aldrich) and 25 µg/ml carbenicillin (carbenicillin disodium salt, Biomol). They were left to dry for one day. Then, L4 larvae were transferred to the plates and were allowed to grow for at least 24 h until the adult stage. The eggs were then used for experiments and were transferred into microchambers for imaging. The bacteria used in the microchambers were the same dsRNA expressing HT115 as the ones on which the parents had been fed.

#### Transcriptome analysis

For transcriptome analysis, we collected 150 L4 larvae during lethargus per condition. We prepared three biological replicates. The larvae were collected in TRIzol (Invitrogen) and stored at -80°C [82]. IMGM Laboratories GmbH performed RNA isolation, RNA sequencing and computation of differential gene expression (DGE) analysis. At IMGM, RNA was isolated using the RNeasy lipid tissue kit (Qiagen) with mechanical homogenization by steel beads and on-column DNase digestion. The RNA concentration and purity were measured using the NanoDrop ND-1000 spectral photometer (Peqlab). The purity was measured by the A260/A280 ratio and only RNA samples with A260/A280 ratios ≥ 1.6 were used for RNA sequencing. RNA degradation was measured on a 2100 Bioanalyzer (Agilent Technologies) using RNA 6000 Pico LabChip kits (Agilent Technologies). The RNA was separated according to fragment size and the RNA integrity number (RIN) calculated from the electrophoretic profile [83]. Only samples with a RIN > 7.5 were used for sequencing. Library preparation was performed with the TruSeq® Stranded mRNA HT technology. 300 ng RNA was fragmented by divalent cations. Using reverse transcription, single cDNA strands were created. For the second cDNA strand, dUTP was incorporated instead of dTTP, to avoid amplification by the DNA polymerase in the subsequent PCR. This guaranteed strandedness. 3’-ends were adenylated and ligated with sequencing primers, flow cell-binding sites, and indices for sequencing of pooled libraries. Adapter-ligated fragments were amplified by PCR. After the PCR and at the end of the library preparation, all samples were quality controlled using DNA 1000 LabChip kits on the 2100 Bioanalyzer. 5.4 ng DNA of each sample was used for final library pooling, subsequently diluted to 2 nM and denatured with NaOH.

Sequencing of the library was performed at a final concentration of 1.8 pM and with a 1 % PhiX v3 control library spike-in (Illumina) on the NextSeq500 sequencing system (Illumina). For cluster generation and sequencing of all samples, 75 cycles of paired-end runs were performed with the final library. Sequencing was controlled by the NextSeq Control Software. Before sequencing, clusters were generated by bridge amplification. The sequencing principle based on the sequencing by synthesis (SBS) approach and two-channel signal detection. By different fluorescent-labeled terminator dNTPs, the nucleotides were determined.

The RNA-Seq analysis was performed with the CLC genomics workbench “RNA-Seq analysis” tool. It maps the next-generation sequencing reads against the current *C. elegans* genome NC_001328 from NCBI. The following mapping parameters were applied: mismatch cost: 2, insertion/deletion cost: 3, length fraction: 0.8, similarity fraction: 0.8, local alignment, maximum number of hits per read: 5.

The expression values were normalized to reads per kilobase of exon model per million mapped reads (RPKM) [84]. The differentially expressed transcripts were calculated. The comparisons were based on the “total exon reads” expression values to allow subsequent statistical analysis by “empirical analysis of DGE”. To analyze significant expression, the exact test for two-group comparisons was used [85]. The dispersion of the data was chosen to be tag wise and the total count filter cutoff threshold was set to 100. To control the false discovery rate (FDR), FDR corrected p-values were calculated and stringent filtering was used. A feature was classified “upregulated” if its FDR-corrected p-value was ≤ 0.05 and fold change was ≥ 2. Analogously, a gene was classified “downregulated” if its FDR-corrected p-value was ≤ 0.05 and fold change was ≤ -2. By using stringent filtering, all features outside of these thresholds were filtered out of the data.

To compare transcriptional changes of *nlp*s and *cnc*s with other transcriptomes, the log fold change was calculated as follows: the log2 of the mean of the total gene reads of all three samples per genotype was calculated and the result of the wild type samples subtracted from the results of the *nas-38(ok3407)* samples. The AMP expression fold change was then compared to the infection transcriptome deriving from RNAseq 12 h after infection [78] and to the fold change between L3 wake and L4 lethargus transcriptome [49].

Tissue enrichment analysis on all genes found upregulated in the *nas-38(ok3407)* transcriptome was performed with the Gene Set Enrichment Analysis tool available from wormbase (Version WS274) using a q value threshold of 0.1.

GO Term analysis was performed using g:profiler. We analyzed the genes found upregulated in the *nas-38(ok3407)* background with default settings. Term size was set between 10 and 1000 to get rid of generic terms.

The heatmap in figure 2A was designed using heatmapper.ca.

#### Epidermal wounding

For laser-induced wounding, we prepared agarose microchambers with young adults. This microchamber was set onto our microscope set-up that was additionally equipped with a 355 nm laser (Rapp Opto, DPSL-355/14, direct coupling). This laser set-up was typically used for laser ablation of cells [86]. The 100x SFluor objective was used and a laser pulse was delivered to the tail or the lateral posterior body. Before laser irradiation, the focal plane was adjusted to 7 µm from the glass surface into the microchamber. The number of laser pulses varied between 1 and 5 and the laser power between 70 % and 100 %, depending on the response behavior. After wounding, the behavior and the neuronal activity were measured. Worms that were strongly harmed and released internal contents were excluded.

For needle-based injury [28, 87], needles were pulled from glass capillaries (GC100F-10, 1mm outer diameter, 0.58 mm inner diameter, Harvard Apparatus) using the Flaming/Brown micropipette puller (Model P-97, Shutter Instruments) with following configuration: Heat: 567, Pull: 110, Vel: 240, Del: 220. Needles were pulled so that they had a stiff pointed end. We wounded worms on the plate without prior immobilization. For needle wounding, only young adults were selected. Under the dissection microscope the worms were pricked at the tail or the lateral posterior body. Worms that lost internal content after wounding were excluded.

In the presence of food, wild type worms showed a highly variable behavioral response to wounding ranging from no sleep to a fraction of 80 % sleep with a median of 14.0 % (Figure S7C). Moreover, worms that were placed in microchambers in the presence of OP50 bacteria often looked sick after a few hours. We hence measured sleep in microchambers without food. Worms were grown on OP50 bacteria, wounded, and then placed in 370 µm x 370 µm x 45 µm agarose microfluidic compartments without food and imaged for 6h directly following the wounding process. Under these conditions, average sleep time was slightly decreased and less variant (Figure 6D). Survival after wounding was scored manually after 24h and 48h on NGM plates with food. The experimenter was blinded to the genotype of the animals during the entire experiment (wounding and scoring).

#### Data availability

FastQ Files and the fold change analyses are available in GEO-NCBI under the accession number GSE146642 and can be viewed at https://www.ncbi.nlm.nih.gov/geo/query/acc.cgi?acc=GSE146642 using the token otchuiucvduprqv.

#### Statistics

Statistical tests used were the two-sample t-test with Welch-correction for suspected unequal variance, the Fishers exact test and the paired Wilcoxon rank test. All tests were performed with MatLab (MathWorks) or OriginPro software. All p-values are FDR-corrected using the Benjamini-Hochberg-correction. The specific tests used are described in the respective figure captions. The boxplots show the 25 % – 75 % percentile, the whiskers indicate the 10 % - 90 % range. The grey line is the median. Fold changes in sleep, lethargus or *nlp-29p::GFP* fluorescence measurements were calculated based on mean values. Larvae that did not develop properly or adult worms that lost internal content after wounding were excluded from all analysis. All plots were designed using OriginPro and Adobe Illustrator software.

---

**Figure S1.**
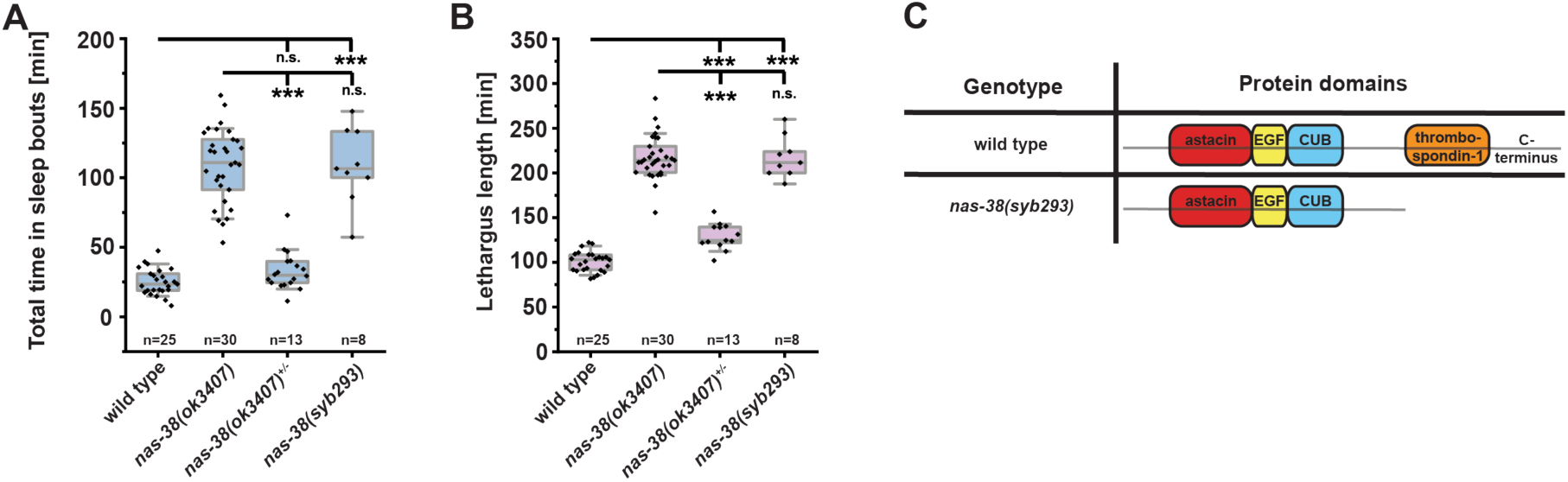
Sleep and lethargus length in *nas-38* truncation mutants. (**A**) Total time spent in sleep of *nas-38* truncation mutants. The CRISPR/Cas9-rebuilt clean truncation of *nas-38(ok3407), nas-38(syb293)*, caused similar sleep amounts as the *nas-38(ok3407)* mutants (334.2 % increase compared to wild type, no significant difference compared to *nas-38(ok3407)*). (**B**) Lethargus lengths of the additional *nas-38* mutants. Heterozygous *nas-38(ok3407)* increased lethargus duration by 28.1 %. The truncated *nas-38(syb293)* allele increases lethargus duration by 116.0 %. *** denotes p ≤ 0.001, * denotes p ≤ 0.05, two sample t-test. The boxplots show individual data points, the box represents the 25 % – 75 % range, the whiskers cover the 10 % - 90 % range, the thin gray line represents the median. (**C**) The protein structure of the *nas-38* c-terminal truncation alleles.

**Figure S2.**
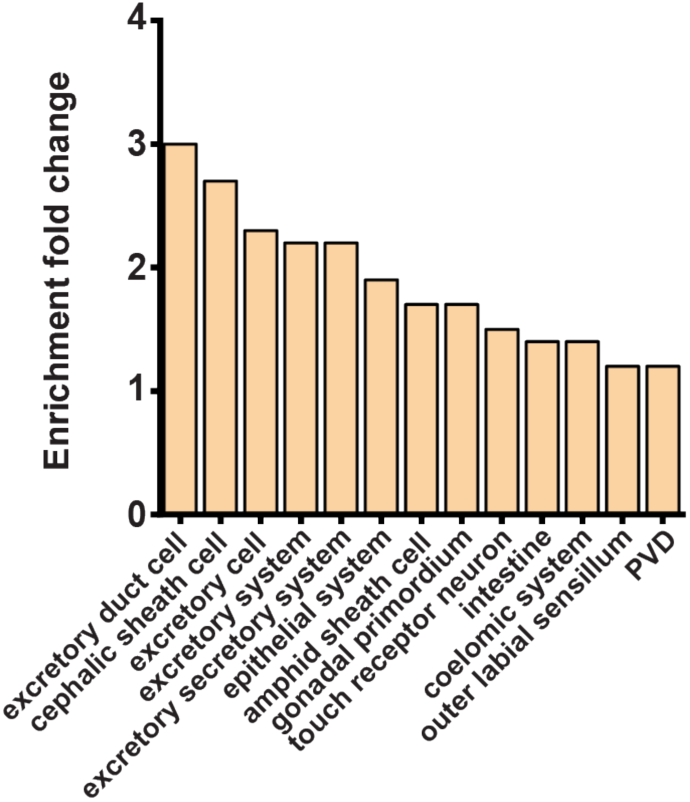
Tissue enrichment analysis of the *nas-38(ok3407)* transcriptome. Differentially expressed genes in *nas-38(ok3407)* are most strongly associated with secretion.

**Figure S3.**
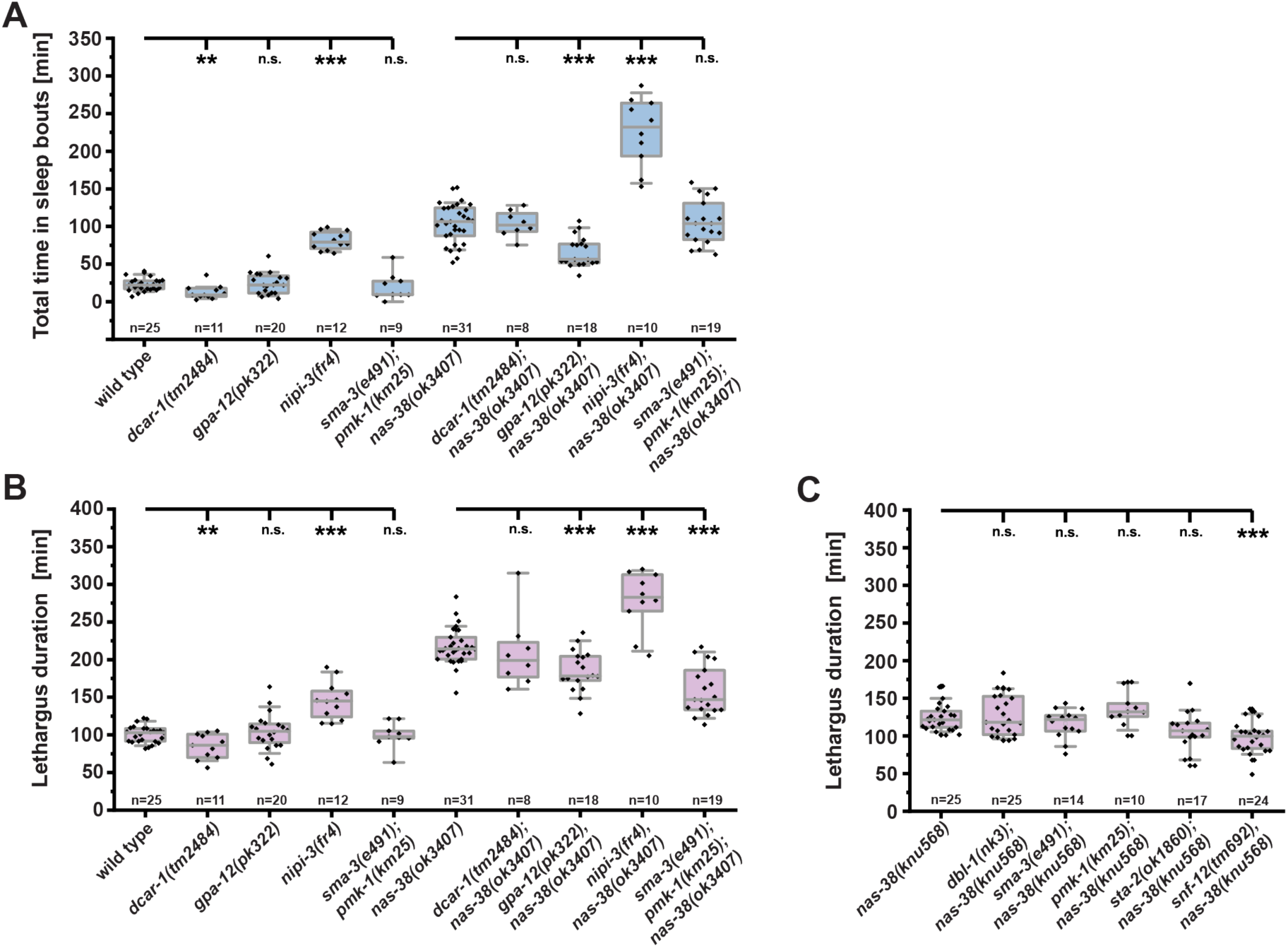
Suppression of increased sleep amount and lethargus length in *nas-38(ok3407)* by mutations in innate immune response genes. (**A**) Quantification of sleep in *nas-38(ok3407)* combined with mutations in innate immune response genes. *dcar-1(tm2484), nipi-3(fr4)* and the *sma-3(e491);pmk-1(km25)* double mutant did not suppress increased sleep caused by *nas-38(ok3407). gpa-12(pk322)* reduced increased sleep caused by *nas-38(ok3407)* by 36.5 %. (**B**) Quantification of lethargus length in *nas-38(ok3407)* combined with mutations in innate immune response genes. *dcar-1(tm2484)* and *nipi-3(fr4)* did not suppress increased lethargus length caused by *nas-38(ok3407). gpa-12(pk322)* decreased lethargus duration by 14.9 %,, and the *sma-3(e491); pmk-1(km25)* double mutant by 27.1 %. (**C**) *snf-12(tm692)* decreased lethargus length in *nas-38(knu568)* by 21.2 %. ** denotes p ≤ 0.01, *** denotes p ≤ 0.001, two sample t-test. The boxplots show individual data points, the box represents the 25 % – 75 % range, the whiskers cover the 10 % - 90 % range, the thin gray line is the median.

**Figure S4.**
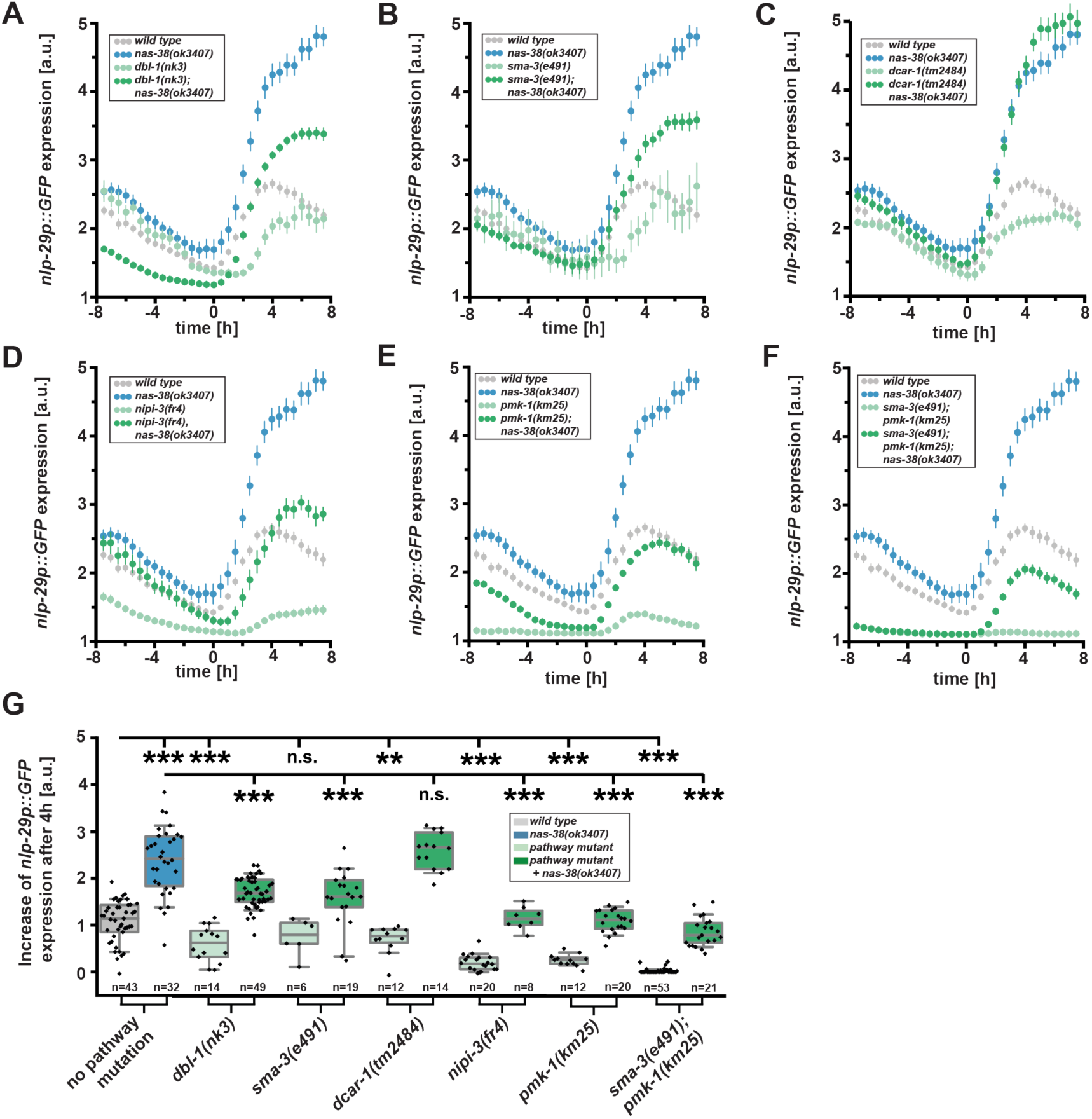
Effects of mutation of innate immune response genes on NLP-29 expression in *nas-38(ok3407)* (**A**) – (**F**) Quantification of *nlp-29p::GFP* intensity ranging from 8 h before to 8 h after lethargus onset in *nas-38(ok3407)* combined with either *dbl-1(nk3), sma-3(e491), dcar-1(tm2484), nipi-3(fr4), pmk-1(km25)* or the *pmk-1(km25);sma-3(e491) double mutant.* Wild type expression levels are shown in grey, expression levels in *nas-38(ok3407)* are shown in blue, expression levels in single mutants in light green and mutant combination with *nas-38(ok3407)* in dark green. Shown are mean values ± SEM. The same wild type and *nas-38(ok3407)* data as in figures 4A-D are shown in every panel. (**G**) *nlp-29p::GFP* expression over the first 4 h after lethargus onset. *dbl-1(nk3)* suppressed *nas-38(ok3407)*-induced *nlp-29p::GFP* expression by 28.5 %, *sma-3(e491)* by 33.4 %, *nipi-3(fr4)* by 51.8 %, *pmk-1(km25)* by 54.2 %, and the *sma-3(e491);pmk1-(km25)* double mutant by 64.0 %. *** denotes p ≤ 0.001, ** denotes p ≤ 0.01, two sample t-test. The boxplots show individual data points, the box represents the 25 % – 75 % range, the whiskers cover the 10 % - 90 % range, the thin gray line is the median.

**Figure S5.**
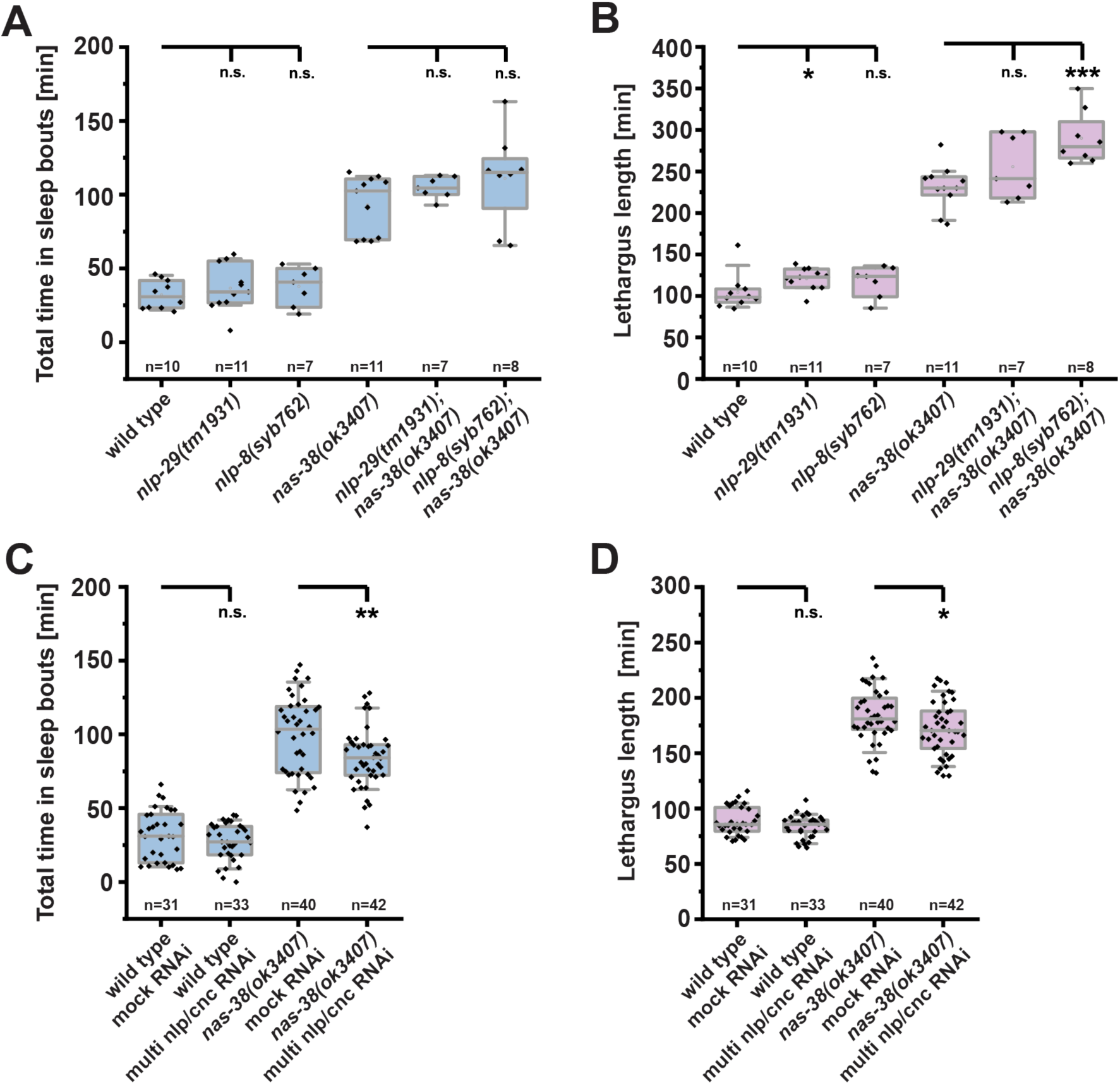
Effects of single *nlp* mutations and of RNAi of multiple *nlp* and *cnc* genes on *nas-38(ok3407)-*induced sleep. (**A**) Quantification of sleep in *nlp-29* and *nlp-8* deletion mutants alone and in combination with *nas-38(ok3407)*. Neither *nlp-29(tm1931)* nor *nlp-8(syb762)* suppressed increased sleep caused by *nas-38(ok3407)*. (**B**) Quantification of lethargus duration in the single *nlp* deletion mutants alone and in combination with *nas-38(ok3407)*. Increased lethargus length caused by *nas-38(ok3407)* was not suppressed by either *nlp-29* or *nlp-8* mutation. (**C**) Quantification of sleep in wild type worms and *nas-38(ok3407)* following RNAi of multiple *cnc* and *nlp* genes. RNAi did not affect wild type sleep amount but reduced sleep in *nas-38(ok3407)* by 15.3 %. (**D**) Quantification of lethargus duration following RNAi of multiple *cnc* and *nlp* genes significantly reduced lethargus length in *nas-38(ok3407)* by 6.4 %. *** denotes p ≤ 0.001, ** denotes p ≤ 0.01, * denotes p ≤ 0.05, two sample t-test. The boxplots show individual data points, the box represents the 25 % – 75 % range, the whiskers cover the 10 % - 90 % range, the thin gray line is the median.

**Figure S6.**
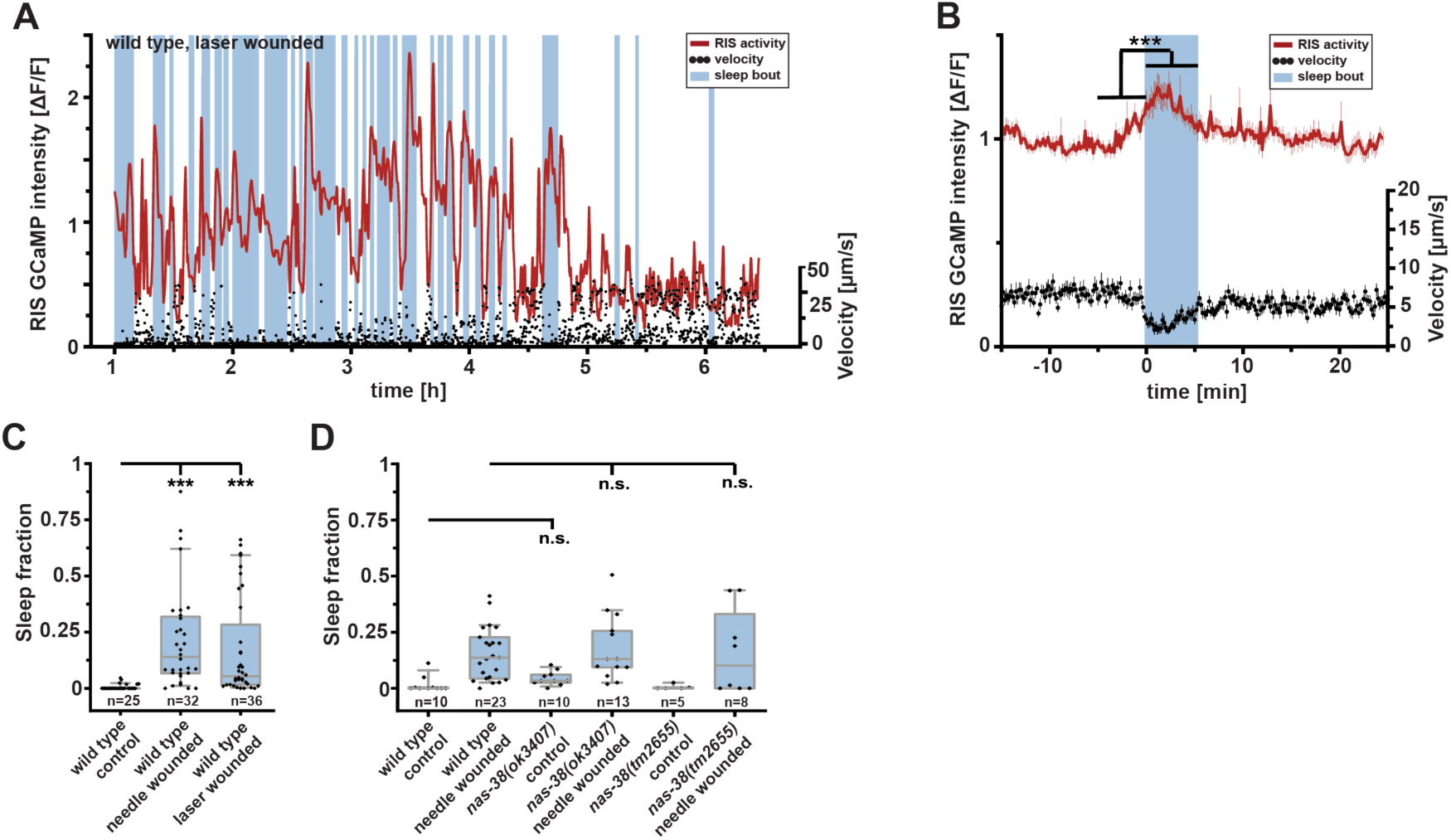
Laser-induced wounding promotes sleep, and sleep after needle-wounding is normal in *nas-38* mutants. (**A**) Velocity data of an individual worm after laser wounding (black) and RIS calcium activity levels (red). Sleep bouts as defined as a cessation of locomotion are labelled in blue. Laser wounding promoted sleep bouts and RIS activated during these sleep bouts. (**B**) Velocity and RIS activity data were averaged by aligning all sleep bout to their onset. RIS activity (ΔF/F) increased by 14.2 % during sleep bouts. n = 29 worms; *** denotes p ≤ 0.001, paired Wilcoxon rank test. (**C**) Quantification of sleep caused by laser wounding and comparison with needle wounding. Average sleep fraction for laser-wounded worms is 17.2 % and for needle-wounded worms it is 21.5 %. Laser-wounded worms showed a highly variable increase in sleep. *** denotes p ≤ 0.001, two sample t-test. (**D**) *nas-38(ok3407)* and *nas-38(tm2655)* mutants had normal sleep behavior both with and without wounding. The wild type reference data are the same as in Figure 6D. The boxplots show individual data points, the box represents the 25 % – 75 % range, the whiskers cover the 10 % - 90 % range, the thin gray line is the median.

**Figure S7.**
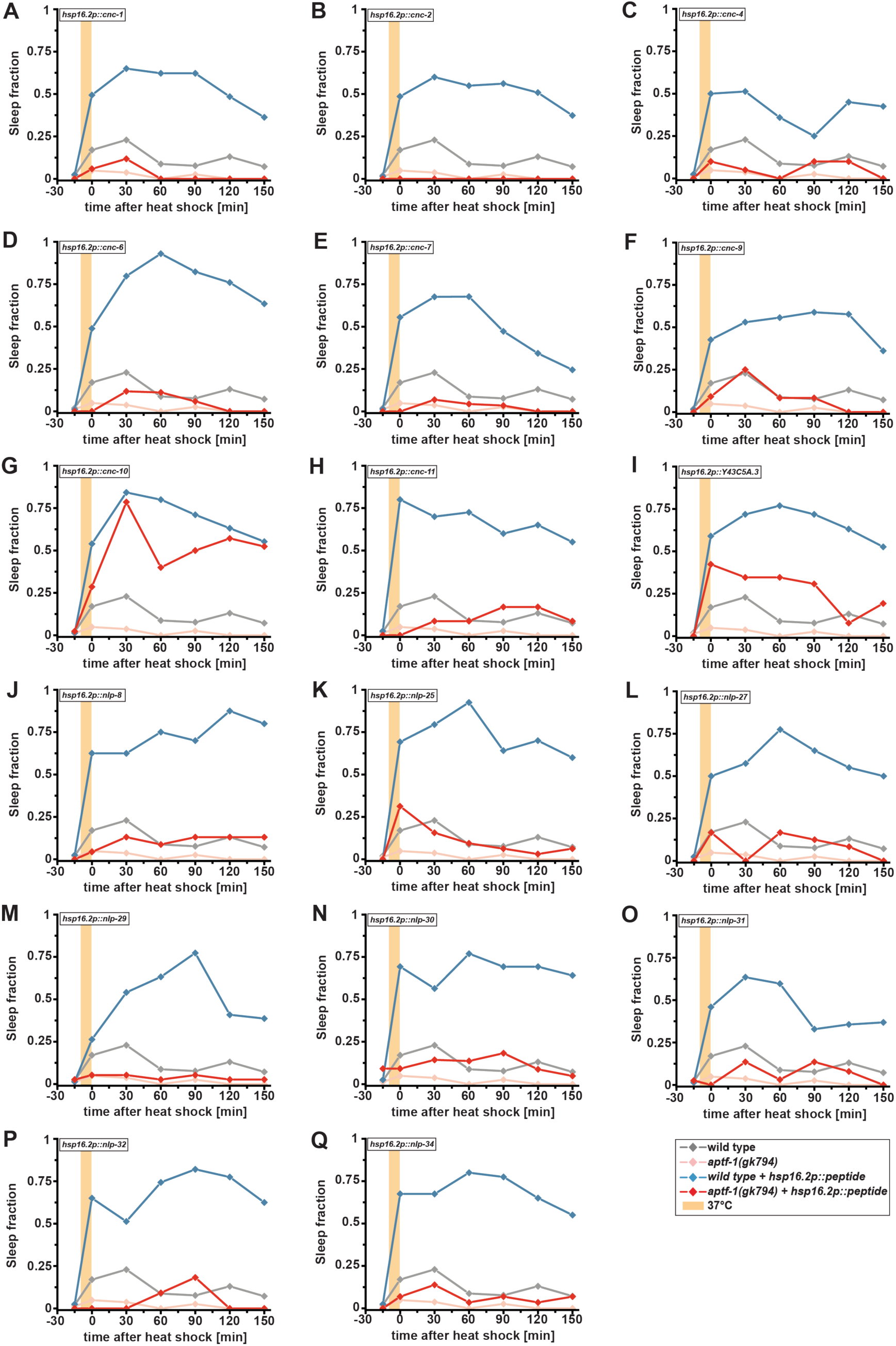
Overexpression of individual *nlp* and *cnc* genes causes behavioral quiescence that depends on RIS. (**A**) - (**Q**) Quantification of behavioral quiescence over time following heat shock-induced overexpression of *nlp* and *cnc* genes in wild type and in *aptf-1(gk794)* mutant backgrounds. The same wild type control (gray) and *aptf-1(gk794)* mutant (light red) measurement is shown in each panel. In each panel the overexpressed (OE)-transgene in wild type background is shown in blue and the respective OE-transgene in the *aptf-1(gk794)* mutant background in red. The 37 °C heat shock is indicated in light orange. Sleep was scored manually by the absence of locomotion and pharyngeal pumping. The first measurement right after the heat shock was defined as t = 0 min.

## References

1. Majde, J.A., and Krueger, J.M. (2005). Links between the innate immune system and sleep. The Journal of allergy and clinical immunology 116, 1188–1198.

2. Zielinski, M.R., and Krueger, J.M. (2011). Sleep and innate immunity. Front Biosci (Schol Ed) 3, 632–642.

3. Toth, L.A., Tolley, E.A., and Krueger, J.M. (1993). Sleep as a prognostic indicator during infectious disease in rabbits. Proc Soc Exp Biol Med 203, 179–192.

4. Bryant, P.A., Trinder, J., and Curtis, N. (2004). Sick and tired: Does sleep have a vital role in the immune system? Nature reviews. Immunology 4, 457–467.

5. Besedovsky, L., Lange, T., and Haack, M. (2019). The Sleep-Immune Crosstalk in Health and Disease. Physiol Rev 99, 1325–1380.

6. Landis, C.A., and Whitney, J.D. (1997). Effects of 72 hours sleep deprivation on wound healing in the rat. Research in nursing & health 20, 259–267.

7. Mostaghimi, L., Obermeyer, W.H., Ballamudi, B., Martinez-Gonzalez, D., and Benca, R.M. (2005). Effects of sleep deprivation on wound healing. Journal of sleep research 14, 213–219.

8. Egydio, F., Tomimori, J., Tufik, S., and Andersen, M.L. (2011). Does sleep deprivation and morphine influence wound healing? Medical hypotheses 77, 353–355.

9. Rechtschaffen, A., and Bergmann, B.M. (1995). Sleep deprivation in the rat by the disk-over-water method. Behavioural brain research 69, 55–63.

10. Saper, C.B., Fuller, P.M., Pedersen, N.P., Lu, J., and Scammell, T.E. (2010). Sleep state switching. Neuron 68, 1023–1042.

11. Weber, F., and Dan, Y. (2016). Circuit-based interrogation of sleep control. Nature 538, 51–59.

12. Bringmann, H. (2018). Sleep-Active Neurons: Conserved Motors of Sleep. Genetics 208, 1279–1289.

13. Krueger, J.M., Frank, M.G., Wisor, J.P., and Roy, S. (2016). Sleep function: Toward elucidating an enigma. Sleep medicine reviews 28, 46–54.

14. Ehlen, J.C., Brager, A.J., Baggs, J., Pinckney, L., Gray, C.L., DeBruyne, J.P., Esser, K.A., Takahashi, J.S., and Paul, K.N. (2017). Bmal1 function in skeletal muscle regulates sleep. eLife 6.

15. Hill, A.J., Mansfield, R., Lopez, J.M., Raizen, D.M., and Van Buskirk, C. (2014). Cellular stress induces a protective sleep-like state in C. elegans. Curr Biol 24, 2399–2405.

16. Van Buskirk, C., and Sternberg, P.W. (2007). Epidermal growth factor signaling induces behavioral quiescence in Caenorhabditis elegans. Nat Neurosci 10, 1300–1307.

17. Boman, H.G. (2003). Antibacterial peptides: basic facts and emerging concepts. J Intern Med 254, 197–215.

18. Toda, H., Williams, J.A., Gulledge, M., and Sehgal, A. (2019). A sleep-inducing gene, nemuri, links sleep and immune function in Drosophila. Science 363, 509–515.

19. Raizen, D.M., Zimmerman, J.E., Maycock, M.H., Ta, U.D., You, Y.J., Sundaram, M.V., and Pack, A.I. (2008). Lethargus is a Caenorhabditis elegans sleep-like state. Nature 451, 569–572.

20. Cassada, R.C., and Russell, R.L. (1975). The dauerlarva, a post-embryonic developmental variant of the nematode Caenorhabditis elegans. Dev Biol 46, 326–342.

21. Iwanir, S., Tramm, N., Nagy, S., Wright, C., Ish, D., and Biron, D. (2013). The microarchitecture of C. elegans behavior during lethargus: homeostatic bout dynamics, a typical body posture, and regulation by a central neuron. sleep 36, 385–395.

22. Turek, M., Lewandrowski, I., and Bringmann, H. (2013). An AP2 transcription factor is required for a sleep-active neuron to induce sleep-like quiescence in C. elegans. Curr Biol 23, 2215–2223.

23. Urmersbach, B., Besseling, J., Spies, J.P., and Bringmann, H. (2016). Automated analysis of sleep control via a single neuron active at sleep onset in C. elegans. Genesis 54, 212–219.

24. Nichols, A.L.A., Eichler, T., Latham, R., and Zimmer, M. (2017). A global brain state underlies C. elegans sleep behavior. Science 356.

25. Turek, M., Besseling, J., Spies, J.P., Konig, S., and Bringmann, H. (2016). Sleep-active neuron specification and sleep induction require FLP-11 neuropeptides to systemically induce sleep. eLife 5.

26. Ermolaeva, M.A., and Schumacher, B. (2014). Insights from the worm: the C. elegans model for innate immunity. Seminars in immunology 26, 303–309.

27. Cohen, L.B., and Troemel, E.R. (2015). Microbial pathogenesis and host defense in the nematode C. elegans. Current opinion in microbiology 23, 94–101.

28. Pujol, N., Cypowyj, S., Ziegler, K., Millet, A., Astrain, A., Goncharov, A., Jin, Y., Chisholm, A.D., and Ewbank, J.J. (2008). Distinct innate immune responses to infection and wounding in the C. elegans epidermis. Curr Biol 18, 481–489.

29. Zugasti, O., Bose, N., Squiban, B., Belougne, J., Kurz, C.L., Schroeder, F.C., Pujol, N., and Ewbank, J.J. (2014). Activation of a G protein-coupled receptor by its endogenous ligand triggers the innate immune response of Caenorhabditis elegans. Nature immunology 15, 833–838.

30. Taffoni, C., Omi, S., Huber, C., Mailfert, S., Fallet, M., Rupprecht, J.F., Ewbank, J.J., and Pujol, N. (2020). Microtubule plus-end dynamics link wound repair to the innate immune response. eLife 9.

31. Belougne, J., Ozerov, I., Caillard, C., Bedu, F., and Ewbank, J.J. (2020). Fabrication of sharp silicon arrays to wound Caenorhabditis elegans. Sci Rep 10, 3581.

32. Ewbank, J.J., and Pujol, N. (2016). Local and long-range activation of innate immunity by infection and damage in C. elegans. Current opinion in immunology 38, 1–7.

33. Kim, D.H., Feinbaum, R., Alloing, G., Emerson, F.E., Garsin, D.A., Inoue, H., Tanaka-Hino, M., Hisamoto, N., Matsumoto, K., Tan, M.W., et al. (2002). A conserved p38 MAP kinase pathway in Caenorhabditis elegans innate immunity. Science 297, 623–626.

34. Nicholas, H.R., and Hodgkin, J. (2004). Responses to infection and possible recognition strategies in the innate immune system of Caenorhabditis elegans. Molecular immunology 41, 479–493.

35. Dierking, K., Polanowska, J., Omi, S., Engelmann, I., Gut, M., Lembo, F., Ewbank, J.J., and Pujol, N. (2011). Unusual regulation of a STAT protein by an SLC6 family transporter in C. elegans epidermal innate immunity. Cell host & microbe 9, 425–435.

36. Ziegler, K., Kurz, C.L., Cypowyj, S., Couillault, C., Pophillat, M., Pujol, N., and Ewbank, J.J. (2009). Antifungal innate immunity in C. elegans: PKCdelta links G protein signaling and a conserved p38 MAPK cascade. Cell host & microbe 5, 341–352.

37. Zugasti, O., and Ewbank, J.J. (2009). Neuroimmune regulation of antimicrobial peptide expression by a noncanonical TGF-beta signaling pathway in Caenorhabditis elegans epidermis. Nature immunology 10, 249–256.

38. Wahl, S.M. (2007). Transforming growth factor-beta: innately bipolar. Current opinion in immunology 19, 55–62.

39. Pujol, N., Zugasti, O., Wong, D., Couillault, C., Kurz, C.L., Schulenburg, H., and Ewbank, J.J. (2008). Anti-fungal innate immunity in C. elegans is enhanced by evolutionary diversification of antimicrobial peptides. PLoS pathogens 4, e1000105.

40. Pujol, N., Davis, P.A., and Ewbank, J.J. (2012). The Origin and Function of Anti- Fungal Peptides in C. elegans: Open Questions. Front Immunol 3, 237.

41. Nathoo, A.N., Moeller, R.A., Westlund, B.A., and Hart, A.C. (2001). Identification of neuropeptide-like protein gene families in Caenorhabditiselegans and other species. Proc Natl Acad Sci U S A 98, 14000–14005.

42. Couillault, C., Pujol, N., Reboul, J., Sabatier, L., Guichou, J.F., Kohara, Y., and Ewbank, J.J. (2004). TLR-independent control of innate immunity in Caenorhabditis elegans by the TIR domain adaptor protein TIR-1, an ortholog of human SARM. Nature immunology 5, 488–494.

43. consortium, C.e.k. (2012). large-scale screening for targeted knockouts in the Caenorhabditis elegans genome. G3 (Bethesda) 2, 1415–1425.

44. Bringmann, H. (2011). Agarose hydrogel microcompartments for imaging sleep- and wake-like behavior and nervous system development in Caenorhabditis elegans larvae. J Neurosci Methods 201, 78–88.

45. Friedland, A.E., Tzur, Y.B., Esvelt, K.M., Colaiacovo, M.P., Church, G.M., and Calarco, J.A. (2013). Heritable genome editing in C. elegans via a CRISPR-Cas9 system. Nat Methods 10, 741–743.

46. Park, J.O., Pan, J., Mohrlen, F., Schupp, M.O., Johnsen, R., Baillie, D.L., Zapf, R., Moerman, D.G., and Hutter, H. (2010). Characterization of the astacin family of metalloproteases in C. elegans. BMC developmental biology 10, 14.

47. Corish, P., and Tyler-Smith, C. (1999). Attenuation of green fluorescent protein half-life in mammalian cells. Protein Eng 12, 1035–1040.

48. Hunt-Newbury, R., Viveiros, R., Johnsen, R., Mah, A., Anastas, D., Fang, L., Halfnight, E., Lee, D., Lin, J., Lorch, A., et al. (2007). High-throughput in vivo analysis of gene expression in Caenorhabditis elegans. PLoS biology 5, e237.

49. Hendriks, G.J., Gaidatzis, D., Aeschimann, F., and Grosshans, H. (2014). Extensive oscillatory gene expression during C. elegans larval development. Molecular cell 53, 380–392.

50. Dodd, W., Tang, L., Lone, J.C., Wimberly, K., Wu, C.W., Consalvo, C., Wright, J.E., Pujol, N., and Choe, K.P. (2018). A Damage Sensor Associated with the Cuticle Coordinates Three Core Environmental Stress Responses in Caenorhabditis elegans. Genetics 208, 1467–1482.

51. Kim, D., Grun, D., and van Oudenaarden, A. (2013). Dampening of expression oscillations by synchronous regulation of a microRNA and its target. Nature genetics 45, 1337–1344.

52. George-Raizen, J.B., Shockley, K.R., Trojanowski, N.F., Lamb, A.L., and Raizen, D.M. (2014). Dynamically-expressed prion-like proteins form a cuticle in the pharynx of Caenorhabditis elegans. Biol Open 3, 1139–1149.

53. Miao, R., Li, M., Zhang, Q., Yang, C., and Wang, X. (2020). An ECM-to-Nucleus Signaling Pathway Activates Lysosomes for C. elegans Larval Development. Developmental cell 52, 21–37 e25.

54. Goetting, D.L., Mansfield, R., Soto, R., and Van Buskirk, C. (2020). Cellular damage, including wounding, drives C. elegans stress-induced sleep. Journal of Neurogenetics.

55. Konietzka, J., Fritz, M., Spiri, S., McWhirter, R., Leha, A., Palumbos, S., Costa, W.S., Oranth, A., Gottschalk, A., Miller, D.M., 3rd, et al. (2020). Epidermal Growth Factor Signaling Promotes Sleep through a Combined Series and Parallel Neural Circuit. Curr Biol 30, 1–16 e13.

56. E, L., Zhou, T., Koh, S., Chuang, M., Sharma, R., Pujol, N., Chisholm, A.D., Eroglu, C., Matsunami, H., and Yan, D. (2018). An Antimicrobial Peptide and Its Neuronal Receptor Regulate Dendrite Degeneration in Aging and Infection. Neuron 97, 125–138 e125.

57. White, J.G., Southgate, E., Thomson, J.N., and Brenner, S. (1986). The structure of the nervous system of the nematode Caenorhabditis elegans. Philos Trans R Soc Lond B Biol Sci 314, 1–340.

58. Maluck, E., Busack, I., Besseling, J., Masurat, F., Turek, M., Busch, K.E., and Bringmann, H. (2020). A wake-active locomotion circuit depolarizes a sleep-active neuron to switch on sleep. PLoS biology 18, e3000361.

59. Taylor, S.R., Santpere, G., Reilly, M., Glenwinkel, L., Poff, A., McWhirter, R., Xu, C., Weinreb, A., Basavaraju, M., Cook, S.J., et al. (2019). Expression profiling of the mature <em>C. elegans</em> nervous system by single-cell RNA-Sequencing. bioRxiv, 737577.

60. Kessler, E., Takahara, K., Biniaminov, L., Brusel, M., and Greenspan, D.S. (1996). Bone morphogenetic protein-1: the type I procollagen C-proteinase. Science 271, 360–362.

61. Hishida, R., Ishihara, T., Kondo, K., and Katsura, I. (1996). hch-1, a gene required for normal hatching and normal migration of a neuroblast in C. elegans, encodes a protein related to TOLLOID and BMP-1. EMBO J 15, 4111–4122.

62. Novelli, J., Ahmed, S., and Hodgkin, J. (2004). Gene interactions in Caenorhabditis elegans define DPY-31 as a candidate procollagen C-proteinase and SQT-3/ROL-4 as its predicted major target. Genetics 168, 1259–1273.

63. Davis, M.W., Birnie, A.J., Chan, A.C., Page, A.P., and Jorgensen, E.M. (2004). A conserved metalloprotease mediates ecdysis in Caenorhabditis elegans. Development 131, 6001–6008.

64. Suzuki, M., Sagoh, N., Iwasaki, H., Inoue, H., and Takahashi, K. (2004). Metalloproteases with EGF, CUB, and thrombospondin-1 domains function in molting of Caenorhabditis elegans. Biological chemistry 385, 565–568.

65. Zhang, Y., Li, W., Li, L., Li, Y., Fu, R., Zhu, Y., Li, J., Zhou, Y., Xiong, S., and Zhang, H. (2015). Structural damage in the C. elegans epidermis causes release of STA-2 and induction of an innate immune response. Immunity 42, 309–320.

66. Rohleder, N., Aringer, M., and Boentert, M. (2012). Role of interleukin-6 in stress, sleep, and fatigue. Annals of the New York Academy of Sciences 1261, 88–96.

67. Brenner, S. (1974). The genetics of Caenorhabditis elegans. Genetics 77, 71–94.

68. Wu, Y., Masurat, F., Preis, J., and Bringmann, H. (2018). Sleep Counteracts Aging Phenotypes to Survive Starvation-Induced Developmental Arrest in C. elegans. Curr Biol 28, 3610–3624 e3618.

69. Merritt, C., and Seydoux, G. (2010). Transgenic solutions for the germline. WormBook, 1–21.

70. Sanger, F., and Coulson, A.R. (1975). A rapid method for determining sequences in DNA by primed synthesis with DNA polymerase. Journal of molecular biology 94, 441–448.

71. Redemann, S., Schloissnig, S., Ernst, S., Pozniakowsky, A., Ayloo, S., Hyman, A.A., and Bringmann, H. (2011). Codon adaptation-based control of protein expression in C. elegans. Nat Methods 8, 250–252.

72. Wilm, T., Demel, P., Koop, H.U., Schnabel, H., and Schnabel, R. (1999). Ballistic transformation of Caenorhabditis elegans. Gene 229, 31–35.

73. Praitis, V., Casey, E., Collar, D., and Austin, J. (2001). Creation of low-copy integrated transgenic lines in Caenorhabditis elegans. Genetics 157, 1217–1226.

74. Vallin, E., Gallagher, J., Granger, L., Martin, E., Belougne, J., Maurizio, J., Duverger, Y., Scaglione, S., Borrel, C., Cortier, E., et al. (2012). A genome-wide collection of Mos1 transposon insertion mutants for the C. elegans research community. PLoS One 7, e30482.

75. Reymann, A.C., Staniscia, F., Erzberger, A., Salbreux, G., and Grill, S.W. (2016). Cortical flow aligns actin filaments to form a furrow. eLife 5.

76. Frokjaer-Jensen, C., Davis, M.W., Hopkins, C.E., Newman, B.J., Thummel, J.M., Olesen, S.P., Grunnet, M., and Jorgensen, E.M. (2008). Single-copy insertion of transgenes in Caenorhabditis elegans. Nature genetics 40, 1375–1383.

77. Stringham, E., Pujol, N., Vandekerckhove, J., and Bogaert, T. (2002). unc-53 controls longitudinal migration in C. elegans. Development 129, 3367–3379.

78. Engelmann, I., Griffon, A., Tichit, L., Montanana-Sanchis, F., Wang, G., Reinke, V., Waterston, R.H., Hillier, L.W., and Ewbank, J.J. (2011). A comprehensive analysis of gene expression changes provoked by bacterial and fungal infection in C. elegans. PLoS One 6, e19055.

79. Stringham, E., Dixon, D., Jones, D., and Candido, E. (1992). Temporal and spatial expression patterns of the small heat shock (hsp16) genes in transgenic Caenorhabditis elegans. Molecular biology of the cell 3, 221–233.

80. Timmons, L., and Fire, A. (2001). Ingestion of bacterially expressed dsRNAs can produce specific and potent genetic interference in Caenorhabditis elegans. Gene 263, 103–112.

81. Kamath, R.S., and Ahringer, J. (2003). Genome-wide RNAi screening in Caenorhabditis elegans. Methods 30, 313–321.

82. Simms, D., Cizdziel, P.E., and Chomczynski, P. (1993). TRIzol: A new reagent for optimal single-step isolation of RNA. Focus 15, 532–535.

83. Schroeder, A., Mueller, O., Stocker, S., Salowsky, R., Leiber, M., Gassmann, M., Lightfoot, S., Menzel, W., Granzow, M., and Ragg, T. (2006). The RIN: an RNA integrity number for assigning integrity values to RNA measurements. BMC molecular biology 7, 3.

84. Mortazavi, A., Williams, B.A., McCue, K., Schaeffer, L., and Wold, B. (2008). Mapping and quantifying mammalian transcriptomes by RNA-Seq. Nature methods 5, 621.

85. Robinson, M.D., and Smyth, G.K. (2007). Small-sample estimation of negative binomial dispersion, with applications to SAGE data. Biostatistics 9, 321–332.

86. Fang-Yen, C., Gabel, C.V., Samuel, A.D., Bargmann, C.I., and Avery, L. (2012). Laser microsurgery in Caenorhabditis elegans. In Methods in cell biology, Volume 107. (Elsevier), pp. 177–206.

87. Xu, S., and Chisholm, A.D. (2014). Methods for skin wounding and assays for wound responses in C. elegans. Journal of visualized experiments: JoVE.

